# Beyond enumeration: Phenotype independent “labeling-capture-release” process enabling precise detection of circulating tumour cells and downstream applications

**DOI:** 10.1101/2024.03.27.586941

**Authors:** Zhiqi Lao, Xiaoxue Ren, Dehua Zhuang, Lingxia Xie, Yucong Zhang, Wei Li, Zhenyou Jiang, Yue Chen, Penghui Li, Liping Tong, Paul K. Chu, Huaiyu Wang

**Author notes:** These authors contributed equally (Zhiqi Lao, Xiaoxue Ren).

## Abstract

Although strategies for circulating tumor cells (CTCs) enrichment have been proposed, the practical effects of clinical CTCs detection are far from satisfactory. Generally, the methodologies for CTCs detection aim at naturally occurring targets, but misdetection/interferences are prevalent due to the diverse phenotypes and subpopulations of CTCs with high heterogeneity. Herein, a CTCs isolation system based on the “labeling-capture-release” process is demonstrated for precise and high-efficient enrichment of CTCs from clinical blood samples. The mechanism which is based on abnormal glyco-metabolism of tumor cells including CTCs can be utilized for the surface decoration of CTCs with artificial azido groups. With the aid of bio-orthogonal plates designed with DBCO- and disulfide groups and exploiting the anti-fouling effects, the cells labeled with azido groups can be captured *via* a copper-free click reaction and released in a non-destructive manner during subsequent disulfide reduction. The technique is demonstrated to label multiple different types of tumor cells with the EpCAM+/- phenotypes and adherent/suspended status, and all the epithelial/interstitial/hybrid phenotypes of CTCs can be separated from clinical blood samples from 25 patients with 10 different cancer types. Moreover, our strategy is superior to the clinically approved CTCs detection system from the perspective of broad-spectrum and accurate recognition of heterogeneous CTCs. The capturing efficiency of this isolation system is over 80% and the release efficiency exceeds 90%. Most of the released CTCs survive with maintained glycolytic activity thus boding well for downstream applications such as drug susceptibility tests using viable CTCs.

## Introduction

Circulating tumor cells (CTCs), which are trace viable tumor cells released into the bloodstream from primary/metastatic tumor lesions, can travel in the circulating system to form new metastatic lesions inside the body^1^. CTCs are the leading cause of tumor metastasis and are even responsible for most of the cancer-associated mortality^2^. Apart from the strong correlation with metastasis, CTCs share associated genetic/molecular information with primary/metastatic tumors and hence, detection of CTCs is of clinical significance in early metastasis diagnosis, progress monitoring, treatment evaluation, therapy development, and so on^3^.

Selective enrichment of CTCs from peripheral blood has been explored and techniques based on various mechanisms have been proposed. At present, CTCs detection techniques can be classified into 3 categories according to the targets and/or mechanisms:

a. CTCs isolation based on the physiological characteristics: Relying on the specific physiological properties of CTCs, techniques including filtration^4–10^, centrifugation^11,12^, and microfluidics^13–25^ have been developed to separate CTCs from blood samples directly. The isolation can enrich CTCs with the complete cellular structure and high viability can be attained with elaborate manipulation. However, the physiological features of CTCs overlap those of normal blood cells to a certain extent, thereby producing serious interference, especially when considering the significant quantity difference between normal blood cells and CTCs (millions of normal blood cells *vs.* few to tens of CTCs per milliliter of blood with red blood cells being excluded).
b. CTCs capturing using surface antigens: There are various tumor-associated antigens and/or tumor-specific antigens (epithelial cell adhesion molecule (EpCAM), folic acid receptor, etc.) on the surface of CTCs that can be utilized as the targets to capture CTCs. The capturing process commonly involves affinitive binding between the surface antigens of CTCs and corresponding ligand molecules (antibodies^26–40^, aptamers^41–45^, etc.) decorated on well-designed materials/devices to provide counting results. However, CTCs capturing based on specific antigen recognition is inevitably accompanied by misdetection of antigen-negative CTCs subpopulations. For example, transference of CTCs usually involves an epithelial-mesenchymal transition (EMT) process to down-regulate the EpCAM expression on CTCs, whilst partial CTCs lose all the surficial EpCAM molecules after undergoing EMT^46^. Consequently, the EpCAM-targeted strategies cannot be applied to capture EpCAM-negative CTCs.
c. CTCs detection by combined strategies: CTCs detection based on single strategies is usually plagued by false-negative/positive interferences. In this regard, double and even triple strategies combining physical screening and antigen recognition have been proposed to obtain CTCs with higher purity^47,48^. However, combined detection is complex operationally and can compromise the viability of captured CTCs and even affect the accuracy of detection, as CTCs are susceptible and errors can be introduced during the time-consuming and multiple steps. Although progress has been made in CTCs detection, and even several products/devices/systems are now commercially available, the lack of a gold standard hampers clinical adoption^49^. Hence, further improving the reliability and precision of CTCs detection is considered as running the last mile of this marathon, which is an urgent mission for optimizing the practical performance.

It is well known that fluctuation of the CTCs phenotypes attributable to tumor heterogeneity is the leading cause of the current drawback plaguing CTCs detection, as different phenotypes of CTCs can co-exist in a cancer patient with dozens of subsets/subpopulations found^50–52^. Therefore, broad coverage of different phenotypes of CTCs is key to solving the problem. Malignant carcinomas generally involve an extra-fast growth stage^53^ because of the abnormally high intracellular metabolic activity shared by most carcinoma cells regardless of the cancer types and/or subpopulations, including highly active CTCs in the bloodstream^54^. In this respect, the large difference in the metabolic activity can be exploited to discern CTCs from normal blood cells, and a precise CTCs enrichment approach can be conceived by means of bio-orthogonal metabolic glyco-engineering (MGE). The bio-orthogonal MGE technique developed by Bertozzi et al. is a versatile tool with simple operation^56^. MGE refers to the process in which chemical groups (azido, bicyclo[6.1.0]non-4-yn-9-yl groups, and so on) are introduced onto cell membranes by the intrinsic glyco-biosynthesis pathways in a manner that does not compromise the integrity and activity of cells. The process is usually combined with the classic bio-orthogonal click reactions to anchor functional groups/motifs/materials with the labeled cells for further investigations and biomedical applications^57–61^ such as tumor imaging and tumor targeting. It has been shown that the specific “artificial targets” can be introduced to the surface of almost all types of tumor cells^62–66^. Therefore, it may be possible to incorporate bio-orthogonal chemical motifs onto CTCs *via* MGE to facilitate precise recognition, capturing, and subsequent release of CTCs.

**Scheme 1:**
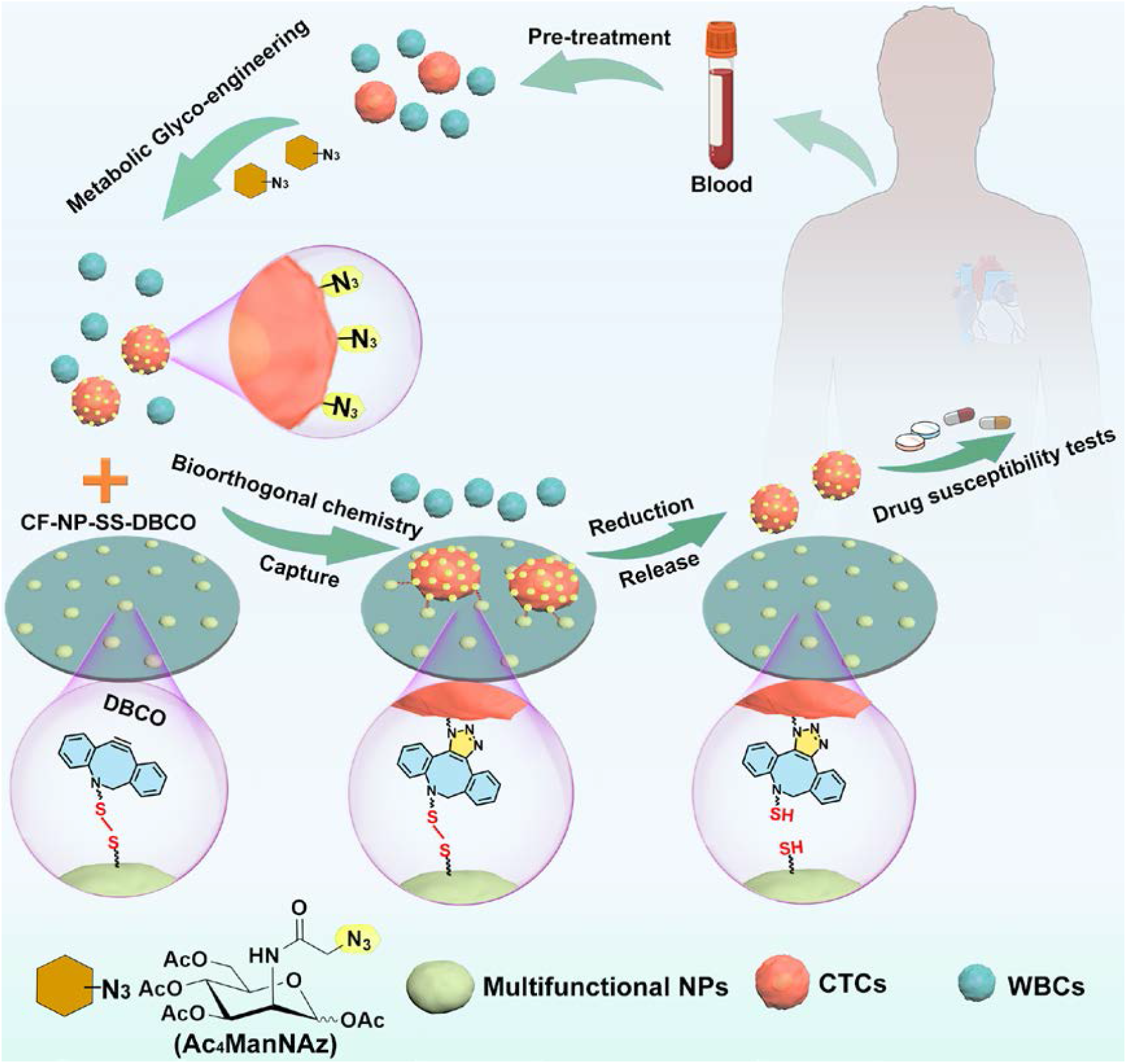
“Labeling-capture-release” process of CTCs based on bio-orthogonal MGE. Herein, a CTCs isolation system based on the “labeling-capture-release” process is described for precise and efficient CTCs enrichment from blood samples followed by downstream applications using the released CTCs. (Scheme 1). The artificial monosaccharide tetra-acetylated *N*-azidoacetyl-D-mannosamine (Ac_4_ManNAz) is used to treat the blood samples, and the surface of rarely existing CTCs can be labeled with azido groups due to the abnormal glyco-metabolism. A capture/release dual-mode plate is prepared by introducing sequentially disulfide and dibenzocyclooctane (DBCO) motifs onto the surface. The functional plate not only captures the azido-labeled cancer cells *via* a copper-free click reaction^67^ between the DBCO and azido groups, but also allows non-destructive release of the captured cells in the ensuing disulfide reduction^68^. This system has the potential to provide a broad-spectrum, direct, and accurate CTCs recognition for various phenotypes despite interferences from normal blood cells. In this study, multiple types of tumor cells with the EpCAM+/- phenotypes and adherent/suspended status are labeled effectively. Using our strategy, all the epithelial/interstitial/hybrid phenotypes of CTCs can be separated from the clinical blood samples of cancer patients. Furthermore, this robust system is credited with the good preservation of cell activity after release and extra low non-specific absorbance of white blood cells (WBCs) under the same conditions, which is highly valuable in the non-invasive cancer diagnosis and can provide great convenience for the downstream various applications of CTCs such as drug susceptibility assessment as preliminarily demonstrated in this study.

## Result

### Optimization of MGE condition

Artificial groups have been conjugated onto viable cells by several intrinsic biosynthetic pathways including the classic Roseman-Warren route^69^, galactosamine (GalN) salvage route^70^, and fucose salvage route^71^. Since both the tumor cells and normal cells can be labeled by MGE and there is a difference between normal blood cells and CTCs in the blood of cancer patients, selective labeling of CTCs from normal blood cells is critical to the isolation strategy. The preferred outcome is that tumor cells are discerned from normal peripheral blood mononuclear cells (PBMCs, chosen as the representative of normal blood cells in this study) by the process of MGE.

To optimize the MGE condition, MCF7 cells, and H524 cells with different biochemical characteristics (EpCAM-positive and EpCAM-negative, adherent and suspended, respectively) are employed to mimic the heterogeneous CTCs. The monosaccharide Ac_4_ManNAz with azido groups are utilized because they participate in intracellular glyco-biosynthesis after co-incubation and so the azido groups are introduced by the Roseman-Warren biosynthetic pathway (Figure 1a). The azido groups on the cells can react with the DBCO-biotin molecules in the copper-free click reaction to immobilize biotin motifs, and the treated cells can be stained with green fluorescence *via* specific binding between biotin and streptavidin-Alexa Fluor 488^®^ (streptavidin-AF488) (Supplementary Figure 1)^70^. Therefore, the fluorescent intensity from the cells after staining is positively related to the amount of azido groups which depends on the amount of added Ac_4_ManNAz and co-incubation time.

**Figure 1.**
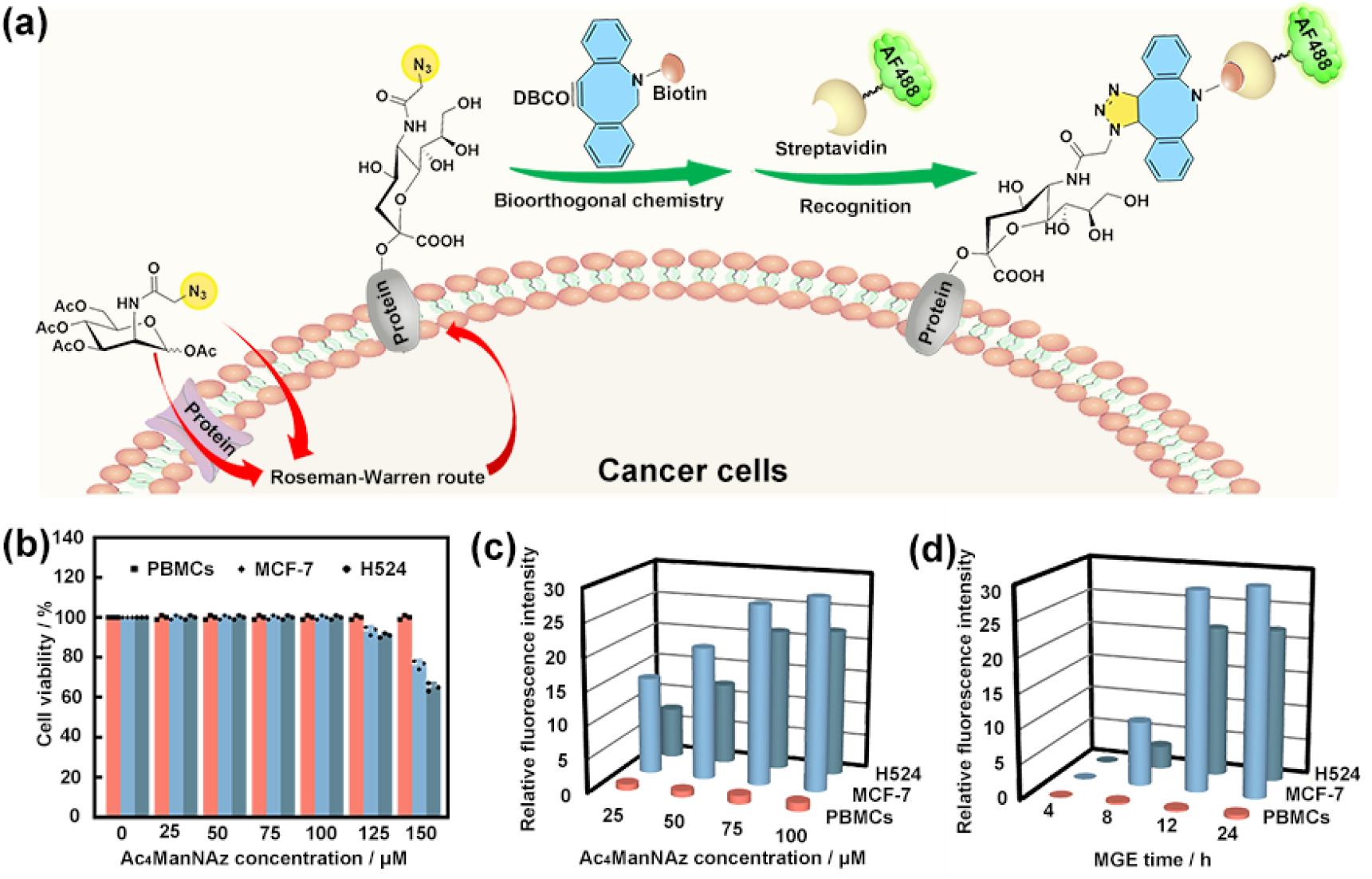
MGE process and fluorescent labeling. **(a)** Schematic diagram showing the introduction of azido groups onto cells *via* the Roseman-Warren route and selective labeling of cells by DBCO-biotin/streptavidin-AF488. (**b)** Monosaccharide concentration-cell survival rate diagram for the determination of the maximum allowable concentration of Ac_4_ManNAz. (**c)** Monosaccharide concentration-cell surface fluorescence intensity diagram explaining the influence of the Ac_4_ManNAz concentration on labeling efficiency. (**d)** MGE time-cell surface fluorescence intensity diagram explaining the influence of the MGE time on labeling efficiency.

According to these points, the cells are first incubated with different concentrations of Ac_4_ManNAz for 24 hours to determine the maximum allowable concentration. As shown in Figure 1b, no obvious cytotoxicity is observed when the MCF7 cells, H524 cells, and PBMCs are cultured with Ac_4_ManNAz at a concentration below 125 μM. In fact, both types of tumor cells are labeled by Ac_4_ManNAz in a concentration-dependent way and the fluorescent intensity from the labeled MCF7 and H524 cells changes only slightly when the concentration of added Ac_4_ManNAz is increased from 75 μM to 100 μM (Figure 1c and Supplementary Figure 2), indicating the azido groups are almost saturated at these concentrations. In contrast, the green fluorescence detected from PBMCs is weak in spite of an Ac_4_ManNAz concentration of 100 μM. Taken together, the optimal concentration of Ac_4_ManNAz is 100 μM.

Since the metabolic activities of CTCs and normal blood cells diminish with time after extraction from blood vessel^73^, the duration of the MGE condition is optimized. Here, the different cells are incubated with 100 μM of Ac_4_ManNAz for 4, 8, 12, and 24 hours prior to fluorescent staining. As shown in Figure 1d and Supplementary Figure 3, both the MCF7 cells and H524 cells show a maximum fluorescent intensity after culturing for 12 hours, whereas fluorescence from the stained PBMCs is near zero under the same conditions. The fluorescent intensity of the labeled cancer cells is at least 40 times higher than that of the labeled PBMCs, implying that tumor cells are MGE-labeled in a specific way. Additional results reveal that 5 other types of tumor cells (A549, Jurkat, HeLa, Huh7, and HepG2 cells) can be labeled under the same conditions (Supplementary Figure 4). Consequently, a phenotype-independent MGE labeling technique (100 μM Ac_4_ManNAz and co-culturing for 12 hours) is employed in the subsequent CTCs isolation study.

### Design, preparation, and characterization of bio-orthogonal films

Capturing of CTCs by bulk biomaterials has been explored and nanostructures with a large surface area are usually prepared on planar biomaterials to enhance the interactions between CTCs and functionalized surfaces^74–83^. Anti-fouling is another requirement for the CTCs-capturing surface and essential to reducing/circumventing nonspecific adsorption of normal blood cells during capturing. Moreover, the functional motifs on the CTCs-capturing surface can be designed to have breakable bonds to facilitate the release of captured CTCs. Based on all these considerations, various surface coatings are designed for the ensuing capturing/releasing study of the azido-labeled CTCs.

In general, 3 DBCO-functionalized chitosan coatings (CF-DBCO, CF-NP-DBCO, and CF-NP-SS-DBCO) are deposited on glass plates for the capturing/releasing study (Figure 2a). Chitosan, a polysaccharide prepared by partial de-acetylation of the -NHAc group on chitin, is chosen as the base material because of the good biocompatibility and provision of multiple free amino groups as reaction sites for conjugation^84–87^. CF-DBCO refers to the chitosan film directly immobilized with the DBCO functional groups, whereas CF-NP-DBCO involves the introduction of DBCO-functionalized nanoparticles onto the chitosan film to enhance the anti-fouling effect^88–91^. The CF-NP-SS-DBCO coating has extra di-sulfide groups compared to CF-NP-DBCO, so that it can be readily reduced to release the captured cells in a mild and non-destructive way.

**Figure 2.**
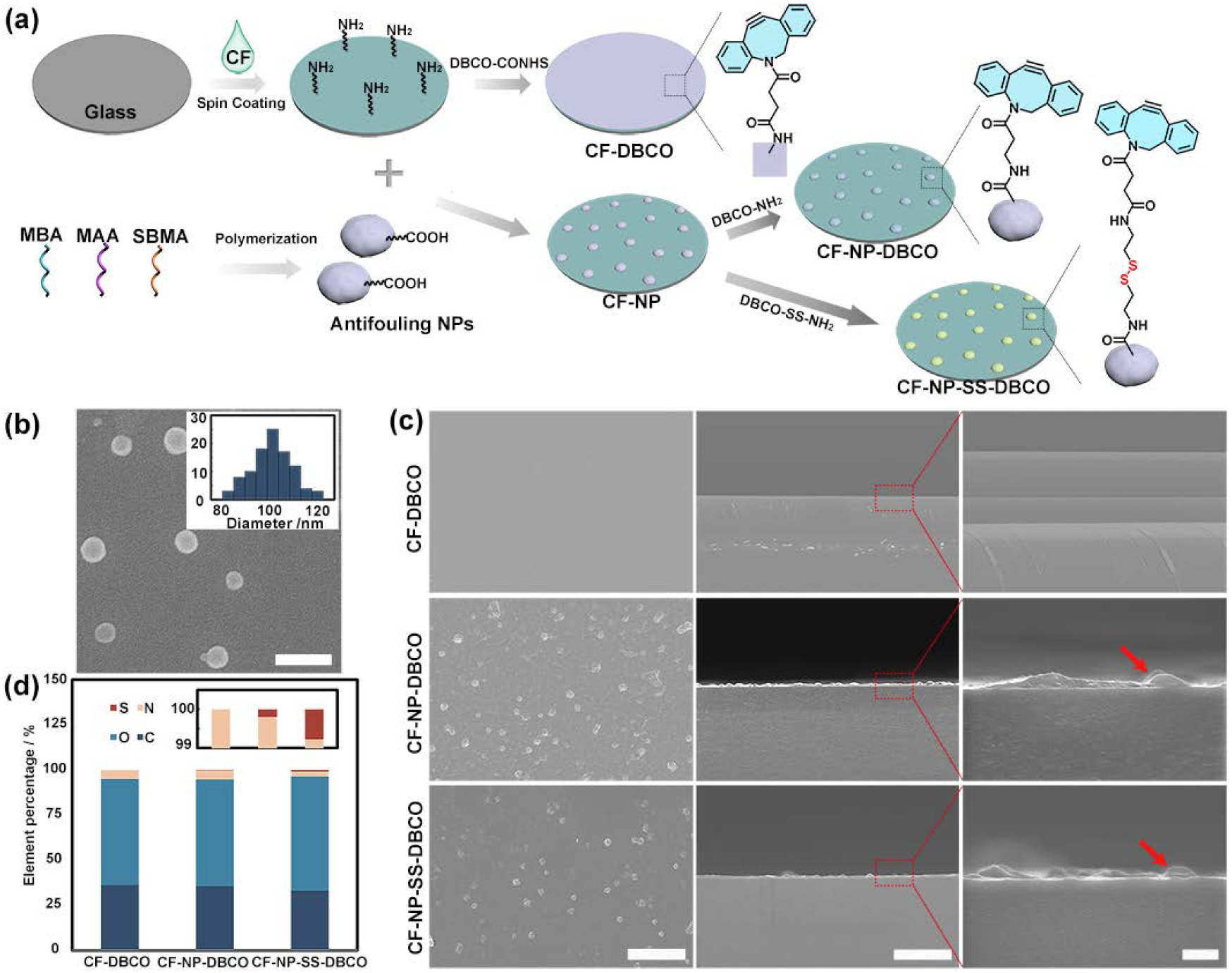
Design, synthesis, and characterization of bio-orthogonal films. **(a)** Schematic diagram showing the preparation of different bio-orthogonal films (Refer to Supplementary Figure 5 for the synthetic details). (**b)** SEM image and size distribution of the anti-fouling nanoparticles (scale bar = 200 nm). (**c)** SEM images of the front and section of different bio-orthogonal films (red arrows indicating embedded nanoparticles) (scale bars = 5 μm, 5 μm, and 500 nm scale bars from left to right, respectively). (**d)** EDS elemental analysis of different bio-orthogonal films.

For the fabrication of CF-DBCO, the chitosan film is deposited on a glass plate by spin-coating followed by treatment with NaOH to expose the amino groups. The chitosan film reacts with DBCO-NHS ester in ethanol to yield CF-DBCO. For the fabrication of the CF-NP-DBCO and CF-NP-SS-DBCO films, sulfobetaine methacrylate (SBMA), methacrylic acid (MAA), and *N, N’*-methylenebisacrylamide (MBA) are adopted for polymerization during refluxing to produce zwitterionic sulfobetaine-type nanoparticles with a diameter of about 100 nm (Figure 2b)^90^. The nanoparticles have abundant carboxylic acid groups which can be immobilized on the chitosan films by the amidation reaction. There are partial carboxylic acid groups on the nanoparticles on the opposite side towards the substrate, and the CF-NP-DBCO and CF-NP-SS-DBCO films are fabricated by subsequent amidation of the remaining carboxylic acid groups with DBCO-NH_2_ and DBCO-SS-NH_2_ groups, respectively (Supplementary Figure 6).

The 3 functionalized samples are characterized by scanning electron microscopy (SEM), energy-dispersive X-ray spectroscopy (EDS), and X-ray photoelectron spectroscopy (XPS). As shown in Figure 2c, the CF-DBCO film consists of 3 layers (DBCO-containing layer, alkalized layer, and acetate layer from top to bottom) with an overall thickness of around 3 μm. The SEM images of CF-NP-DBCO and CF-NP-SS-DBCO substrates reveal that the anti-fouling nanoparticles are dispersed on the films and spread like fried eggs with a diameter of about 400 nm. EDS (Figure 2d) and XPS (Supplementary Figure 7) verify that DBCO motifs exist on the films and sulfur is apparent on CF-NP-SS-DBCO.

### Selective capture and release of cancer cells

After determining the MGE conditions and preparation conditions of different DBCO-functionalized films, CTCs capturing/releasing is assessed (Figure 3a). As MGE is phenotype-independent, H524 cells that resemble the suspension status of CTCs in circulation are chosen based on the optimized MGE conditions (culturing with 100 μM Ac_4_ManNAz for 12 hours). It is presumed that the labeled cells can be captured by the functionalized films *via* a strain-promoted azide-alkyne cycloaddition (SPAAC) reaction between the azido groups on the tumor cells and DBCO groups immobilized on the films. For a comparative study of the azido-positive and azido-negative cells, the H524 cells are treated with Ac_4_ManNAz (azido-positive monosaccharide) and Ac_4_ManNAc (azido-negative monosaccharide), respectively, because both of them can be taken up by cancer cells and metabolized. After MGE labeling, the treated cells are incubated with the 3 DBCO-functionalized films to allow CTCs capture in a spontaneous and time-dependent way. As shown in Figure 3b and Supplementary Figure 8, about 80% of the azido-labeled H524 cells (∼40000 per 50000 cells) are captured by the DBCO-functionalized films within 1 hour, and prolonging the capturing time makes little difference for all 3 films. This stems from surface saturation, and the capacity is more than enough for CTCs detection as the amounts of CTCs in typical clinical samples are quite low (several to dozen CTCs per milliliter of blood). Although the CTCs capturing capability of the different samples is similar, the anti-fouling effects of the CF-DBCO film are inferior to those of the nanoparticle-decorated ones. A small portion of the azido-negative H524 cells absorbs on the CF-DBCO film, whereas almost no non-specific absorption is observed from the CF-NP-DBCO and CF-NP-SS-DBCO films (Figure 3b and Supplementary Figure 8).

**Figure 3.**
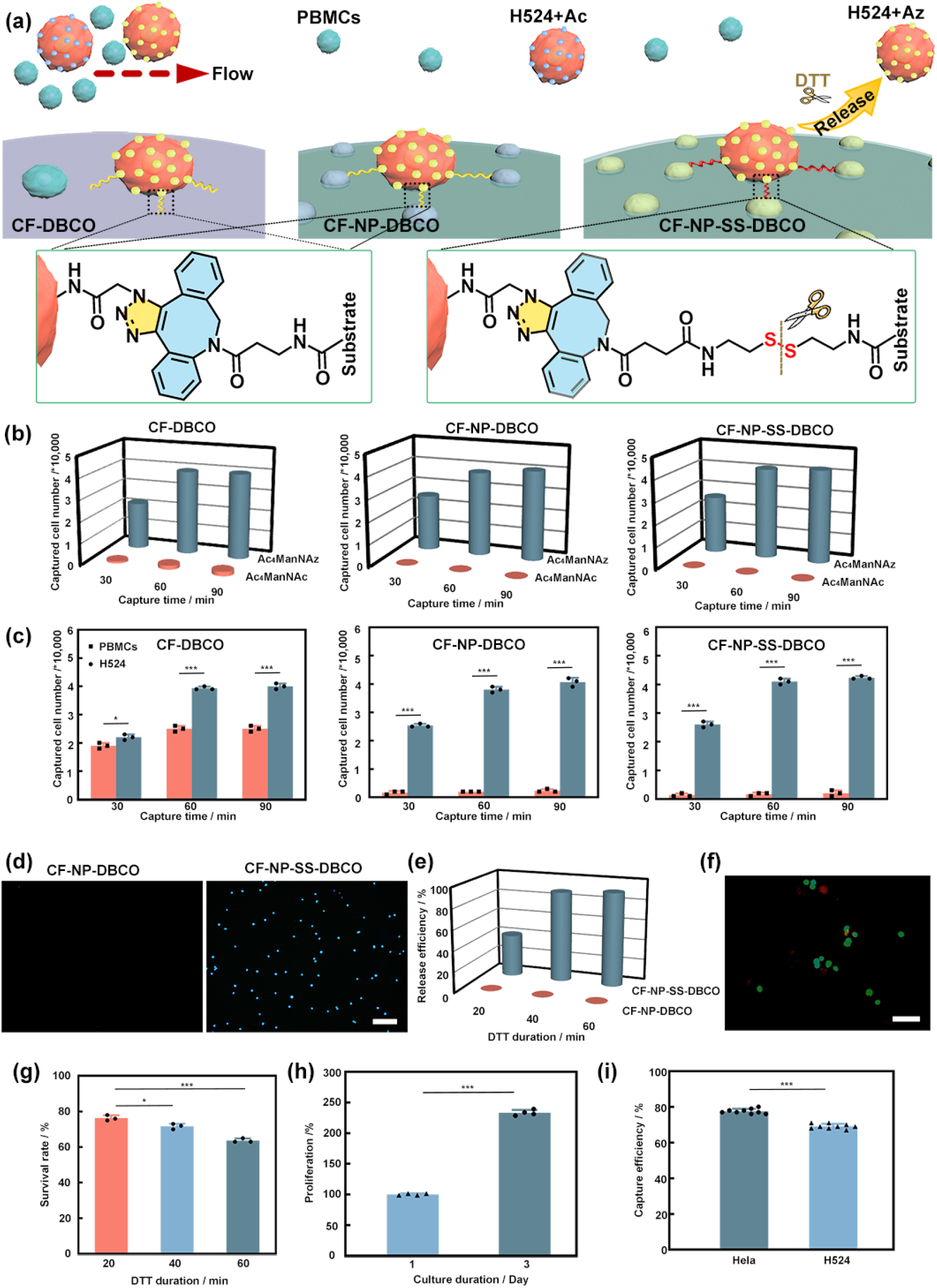
Selective capture and release of cancer cells by bio-orthogonal films. **(a)** Schematic diagram of the capturing and releasing process of cancer cells by the bio-orthogonal films. **(b)** Capture time-captured cell number (from a total of 50000 cells) of MGE-treated H524 cells with/without azido groups on the bio-orthogonal films. **(c)** Capture time-captured cell number (from ∼50000 H524 cells or PBMCs from 1 mL of healthy blood (4∼8 million cells)) of MGE-treated PBMCs and H524 cells on the bio-orthogonal films. **(d)** Fluorescent images of H524 cells (labeled by DAPI) released from the CF-NP-DBCO and CF-NP-SS-DBCO films (scale bar = 100 μm). (**e)** DTT treatment duration-cell release efficiency of CF-NP-DBCO and CF-NP-SS-DBCO. **(f)** Live/dead staining image (green for live/red for dead) of H524 cells released from CF-NP-SS-DBCO after 40 minutes of DTT treatment (scale bar = 50 μm). **(g)** DTT treatment duration-cell survival rates of CF-NP-SS-DBCO. **(h)** Long-term viability of H524 cells after capture and release. **(i)** Artificial CTCs capture efficiency of CF-NP-DBCO.

To further corroborate the biological properties, tests are carried out with both PBMCs and H524 cells. The H524 cells (∼50000 cells) and PBMCs from 1 mL of healthy blood (4∼8 million cells) are first subjected to standard MGE labeling. Under these conditions, the CF-DBCO film exhibits severe non-specific absorption of PBMCs without azido groups, but on the other hand, only a trace amount of PBMCs are observed from the CF-NP-DBCO and CF-NP-SS-DBCO films (Figure 3c), thus confirming the excellent anti-fouling effects. What is more, this strategy is successful in capturing the cancer cells with multiple sizes (Supplementary Figure 9). In comparison with CF-NP-DBCO, CF-NP-SS-DBCO has extra disulfide groups and a study is performed on the release ability with dithiothreitol (DTT) reducing the disulfide bonds. As shown in Figure 3d-e and Supplementary Figure 10, only the CTCs captured by CF-NP-SS-DBCO are released after adding DTT. Release of CTCs is also time-dependent with more than 90% of the captured cells being released in 40 minutes. In addition, most of the released cells remain alive after 40 minutes of DTT treatment (Figures 3f and 3g), of which the long-term viability is even comparable to that of normal cells (Figures 3h and Supplementary Figure 11). By considering the capturing efficiency and anti-fouling effects, the CF-NP-DBCO and CF-NP-SS-DBCO films are more preferred than the CF-DBCO substrate for trace detection of CTCs.

### Artificial CTCs detection using spike-in blood samples

To further verify the feasibility and reliability of our system, a straight-forward simulated experiment of artificial CTCs detection is designed and performed. The artificial CTCs are prepared by spiking HeLa cells or H524 cells (∼100 cells) into 1 mL of fresh blood samples (containing 4∼8 million PBMCs) from healthy donors, and then the mixtures are pre-treated with ACK lysis buffer to remove red blood cells. The remaining cells are collected and subjected to a standard MGE labeling process in the serum-free lymphocyte culture medium. After labeling, the mixed samples are incubated on the CF-NP-DBCO film to allow the capture of artificial CTCs. The captured cells are examined by immunofluorescent staining using nucleus/CD45/pan-CK dyes, and most of them share the feature of tumor cells as nucleus+/CD45-/pan-CK+ (Supplementary Figure 12). It indicates that the artificial CTCs can be selectively labeled by MGE and subsequently captured by the bio-orthogonal films, even in the presence of millions of PBMCs. According to the statistical results, nearly 80% of the HeLa cells and 70% of H524 cells introduced to the blood samples can be captured by this strategy (Figure 3i). It is especially encouraging that the background PBMCs show little interference in the labeling and capturing process.

### Selective labeling of CTCs by MGE

As aforementioned, metabolic labeling trace CTCs with different phenotypes from millions of normal WBCs are required for clinical adoption. Therefore, the feasibility of MGE labeling is evaluated after CTCs are separated from 4 samples with 3 different cancer types by using commercial ISET products (Figure 4a). Here, the whole blood sample (5 mL) drawn from each cancer patient is pre-treated with the ACK lysis buffer to remove red blood cells and then filtered with membranes with standard 8 μm diameter holes to obtain CTCs. The CTCs and some WBCs contaminants are collected and processed under the standard MGE conditions in the serum-free lymphocyte culture medium. Identification of CTCs is performed by immunofluorescent analysis with nucleus/CD45/pan-CK/azido or nucleus/CD45/EpCAM-CKs/azido fluorescent staining. CTCs are identified as nucleus+/CD45-together with at least 1 clinical CTCs biomarker positive (pan-CK or EpCAM-CKs) and WBCs as nucleus+/CD45+ with clinical CTCs biomarker being negative.

**Figure 4.**
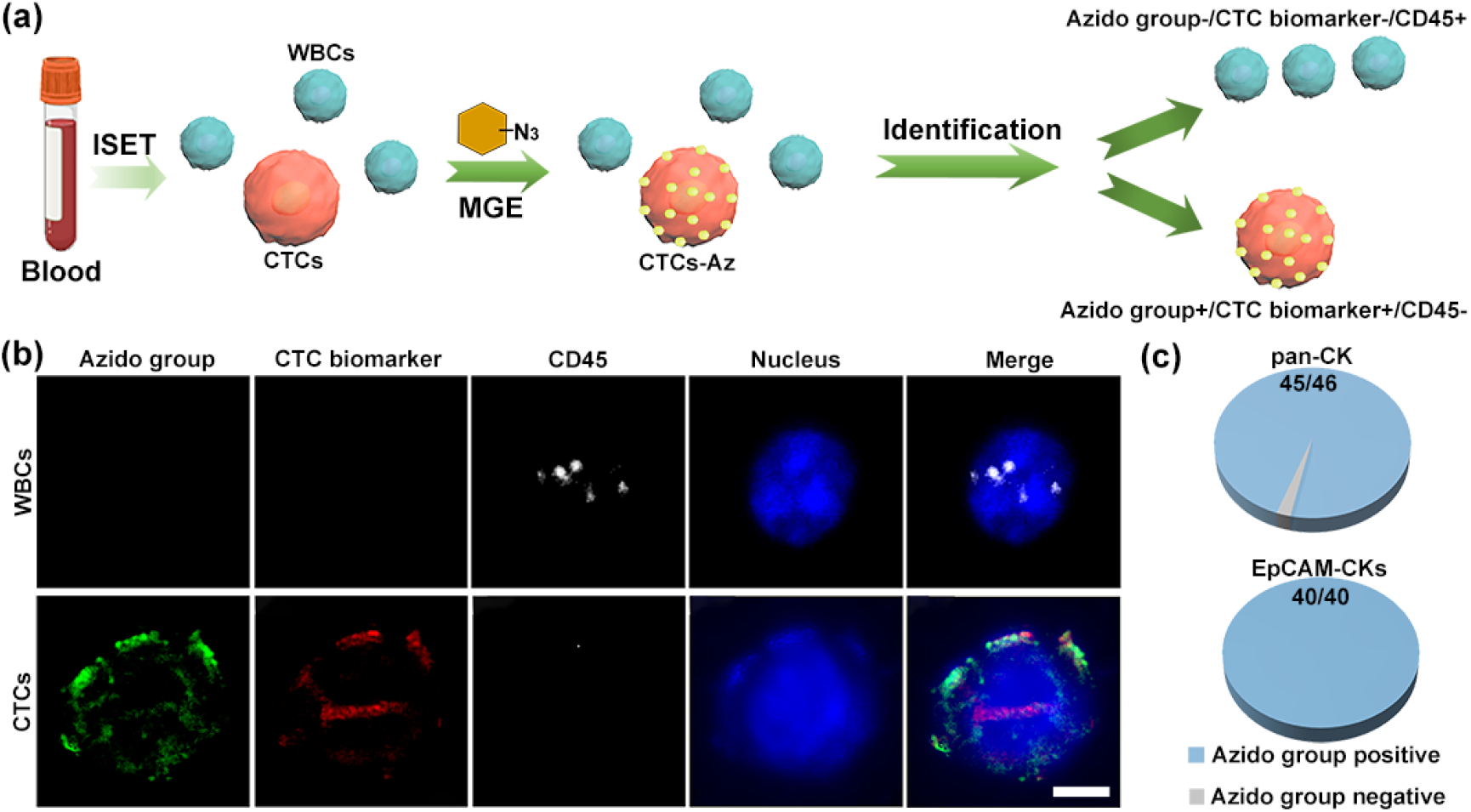
Selective labeling of CTCs in clinical samples by MGE. **(a)** Flow chart of clinical CTCs isolation by ISET, MGE treating process, and immunofluorescent analysis with a combination of nucleus/CD45/CTC marker/azido fluorescent staining (pan-CK represents a mixture of CK1, CK3-8, CK10, CK13-19; EpCAM-CKs represents a mixture of EpCAM, CK8, CK18, CK19). **(b)** Fluorescent images of WBCs and CTCs identified by immunofluorescent analysis (scale bar = 5 μm). **(c)** Statistical analysis of the azido-labeled CTCs in the blood samples.

As shown in Figure 4b, the CTCs that are nucleus+/CD45-/pan-CK+ or nucleus+/CD45-/EpCAM-CKs+ can be stained by the DBCO-biotin/streptavidin-AF488 system (green fluorescence), revealing the good consistency between clinical staining of CTCs and our azido labeling strategy. Among the 46 CTCs identified by the pan-CK marker, 45 of them are successfully labeled with azido groups, and all of the 40 CTCs identified by the EpCAM-CKs marker are labeled with azido groups (Figure 4c, Supplementary Figures 13 and 14). The results demonstrate the feasibility of selective labeling CTCs in clinical samples by MGE and suggest that the artificial groups introduced to the CTCs *via* MGE can be used as general neo-markers for CTCs detection.

### Phenotype-independent capture of CTCs by bio-orthogonal films

To investigate the clinical applicability of our detection system, 1 mL of whole blood drawn from 8 randomly selected cancer patients (P1 to P8) and 2 healthy people (H1 and H2) (For detailed information about donors, please see Supplementary Table 1) is pre-treated with the ACK lysis buffer to remove red blood cells and then undergoes standard MGE labeling in the serum-free lymphocyte culture medium, respectively. Afterward, the treated cell mixtures are incubated on the bio-orthogonal films to allow CTCs capturing *via* the SPAAC reaction. As the results of azido group labeling show correlation with those of pan-CK/EpCAM-CKs staining of CTCs, cross-validation is performed to classify the subsets of captured CTCs. The cells on the bio-orthogonal surface are analyzed by immunofluorescent analysis with nucleus/CD45/EpCAM/vimentin staining, in which 3 general CTC subsets as epithelial CTCs (nucleus+/CD45-/EpCAM+/vimentin-), interstitial CTCs (nucleus+/CD45-/EpCAM-/vimentin+), and hybrid CTCs (nucleus+/CD45-/EpCAM+/vimentin+) can be classified. In EpCAM and vimentin staining, fluorescence of ≤ 7 is regarded as a negative result^93,94^. As shown in Figure 5b, all the epithelial, interstitial, and hybrid CTCs can be captured by the bio-orthogonal system (typical epithelial CTC, interstitial CTC and hybrid CTC shown in the upper, medium and lower column, respectively). It indicates that the introduction of neo-markers (azido groups) enables precise detection of heterogeneous CTCs in the clinical samples. Based on the immunofluorescent staining results displayed in Supplementary Figure 15, the number of CTCs in different blood samples is counted as shown in Figure 5c. It is clear that all the blood samples from different cancer patients (P1-P8) test positive for CTCs, while those from healthy people (H1 and H2) show negative CTCs results. Another merit of our detection system is that the non-specific absorption of WBCs on the films is small on account of the anti-fouling effects. Compared to the previously reported methods for CTCs detection, the background interference of WBCs is at the same level when using the bio-orthogonal films (Supplementary Table 2).

**Figure 5.**
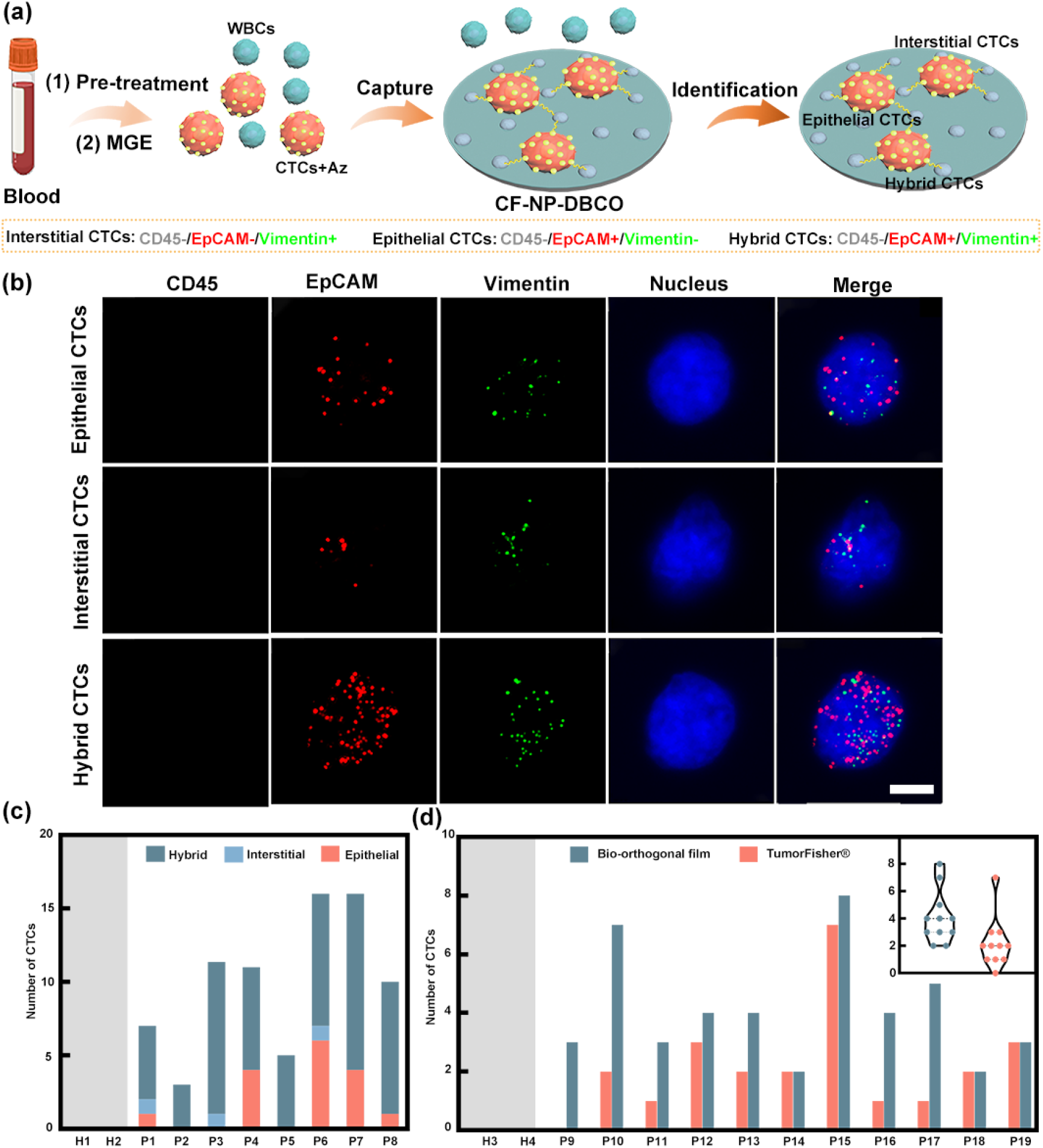
Capturing of CTCs from blood samples by bio-orthogonal films. **(a)** Flow chart illustrating the use of CF-NP-DBCO for CTCs detection. **(b)** General view of the phenotypes of captured CTCs identified by immunohistochemical staining (scale bar = 5 μm). **(c)** Statistical chart of CTCs with different phenotypes captured by CF-NP-DBCO. **(d)** Direct comparison of CTCs detection between CF-NP-DBCO and the TumorFisher® system.

Encouraged by these results, a direct comparison is carried out between our method and a clinically approved CTCs detection system named TumorFisher® (capturing CTCs *via* the EpCAM recognition). The blood samples from another 11 patients with different cancers (P9 to P19) and 2 healthy donors (H3 and H4) are utilized. Each blood sample is halved and parallelly processed by TumorFisher® and our detection workflow, and then the captured CTCs are identified by immunofluorescent staining. As shown in Figure 5d and Supplementary Figure 16, in addition to the negative detection results of healthy people (H3 and H4), our strategy is successful in the trace detection of CTCs from all 11 blood samples from cancer patients (P9-P19). In contrast, 10 of 11 cancer blood samples show positive results, but P9 with colon cancer (partly mesenchymal tissue tumor) is mis-detected (giving a false-negative result) by the commercial TumorFisher® system. Our strategy not only has significant advantages over the TumorFisher® system for the clinical CTCs detection of mesenchymal tissue tumors (P9: 3 vs 0, P10: 7 vs 2, P11: 3 vs 1), but also shows superior detection capability when other kinds of cancers are involved (P12-P19: 5 wins, 3 draws) (Supplementary Figure 16). All in all, the MGE strategy proposed in our study has great potential in the precise labeling of CTCs regardless of phenotypes and subsets.

### Release of viable CTCs for drug susceptibility test

As aforementioned, the CF-NP-SS-DBCO films designed with disulfide groups enable the release of viable cells by subsequent disulfide reduction. Therefore, the whole process of CTCs “labeling-capture-release” is performed, and downstream applications of released CTCs are assessed by testing their susceptibility to different drugs *via* glycolytic activity evaluation (Figure 6a)^96^. In particular, the blood samples from 5 patients (P20 to P24) with stage III colon cancers (For detailed information about donors, please see Supplementary Table 1) are utilized and the captured and subsequently released CTCs are discerned as Nucleus+/CD45-. Meanwhile, 2-Deoxy-2-[(7-nitro-2,1,3-benzoxadiazol-4-yl)amino]-D-glucose (2-NBDG) with green fluorescence is used to identify the glycolytic activity of released CTCs as shown by the fluorescent image in Figure 6b.

**Figure 6.**
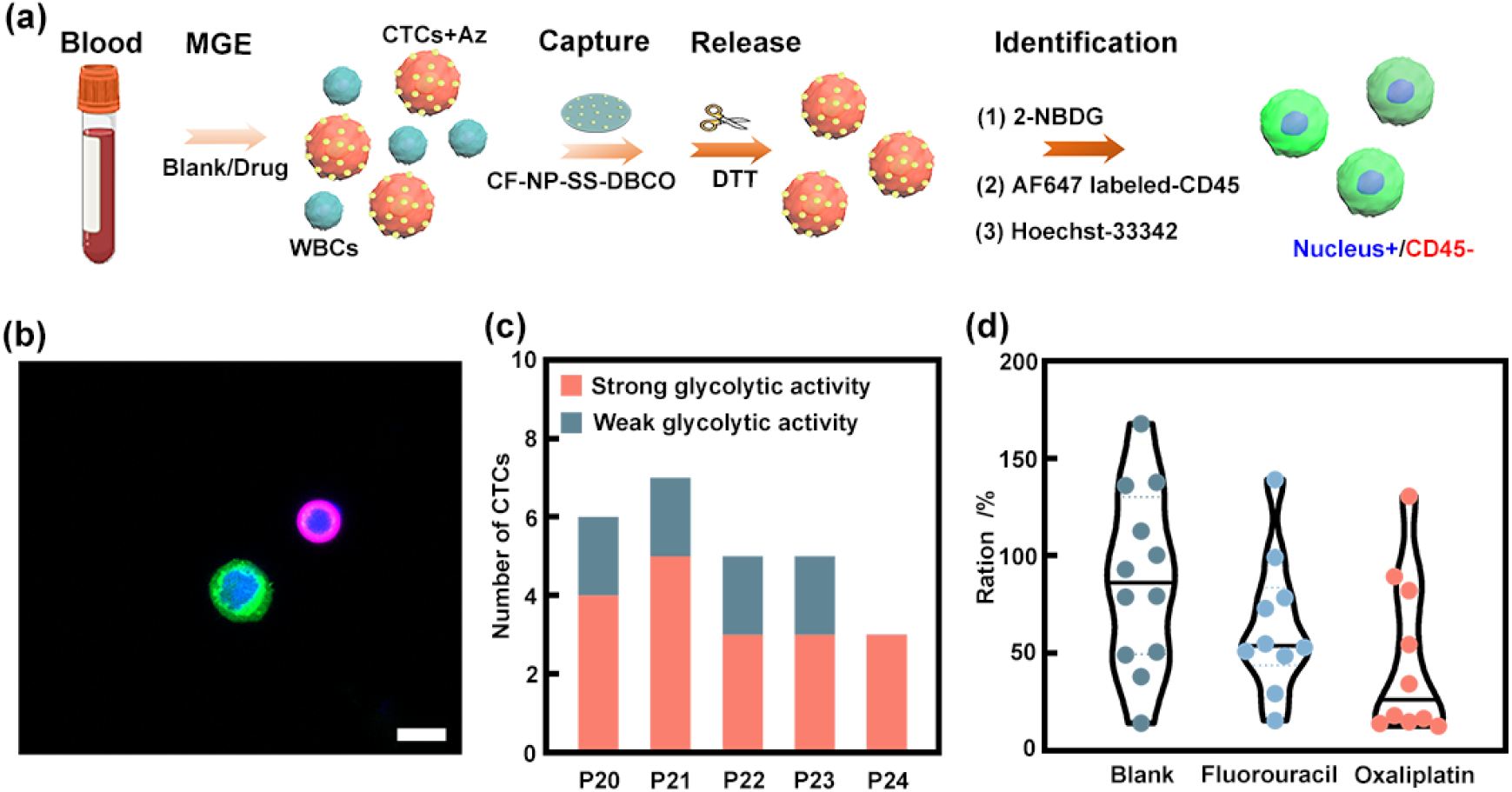
Glycolytic activity of released CTCs and drug susceptibility test. **(a)** Flow chart illustrating the release of CTCs from bio-orthogonal films and tested for the glycolytic activity. **(b)** General view of the released CTCs and WBCs identified by immunohistochemical staining (P20, scale bar = 10 μm). **(c)** Statistical chart of the glycolytic activity of released CTCs (P20-P24). **(d)** Drug susceptibility test using viable CTCs (P25).

The results are very encouraging in that most of the CTCs (Nucleus+/CD45-) after the “labeling-capture-release” process are viable and exhibit good glycolytic activity (Figure 6c and Supplementary Figure 17). Subsequently, the viable CTCs obtained from another patient (P25) with stage III colon cancer (For detailed information about donors, please see Supplementary Table 1) are evenly divided into three portions and tested for their drug susceptibility to Fluorouracil or Oxaliplatin (2 first-line chemotherapeutic drugs for the treatment of stage III colon cancers^97^). As shown in Figure 6d and Supplementary Figure 18, both Fluorouracil and Oxaliplatin interventions can down-regulate the glycolytic activity of tested CTCs, and the Oxaliplatin group delivers the best outcome, affirming that this cancer patient is more susceptible to Oxaliplatin than Fluorouracil. In addition to the drug susceptibility test, other downstream applications of the released CTCs with good viability can be foreseen.

## Discussion

After decades of continuous exploration, progress has been made in CTCs detection. The Cellsearch system was the first FDA-approved product targeting the EpCAM antigens on CTCs. The ISET membrane has been developed to obtain CTCs with intact morphology by physical filtration. Some microfluidic systems have also been designed for direct enrichment of CTCs in blood samples with a high throughput^14–17^. Nowadays, novel CTCs detection techniques with combined mechanisms are emerging but nevertheless, a deeper understanding of the high heterogeneity of CTCs requires more precise and efficient isolation, and the recognition of rare CTCs from millions of WBCs is vital as well. In this respect, the high cellular activity of CTCs originating from abnormal intracellular metabolism provides a promising recognition target. Metastasis begins in the very early stage of cancer, while most of the tumor cells released from lesions die of shear stress, anoikis or immune attack, and so on, and consequently, only a small fraction of highly active tumor cells can survive and turn into CTCs. The high activity is shared by all CTCs regardless of phenotypes, subsets, and subpopulations.

In this study, Ac_4_ManNAz, a monosaccharide used in labeling dozens of different tumor cells by MGE is demonstrated to be able to distinguish CTCs from WBCs by the optimized MGE labeling process. By adjusting the MGE conditions, artificial neo-markers (azido group) can be selectively introduced onto CTCs without compromising their integrity. The condensed neo-markers serve as recognition tags for subsequent capture of CTCs. Although our study demonstrates the great potential of MGE for precise labeling of CTCs, it should be noted that the Ac_4_ManNAz monosaccharide metabolized by the Roseman-Warren route is only one of the choices, and future research on CTCs capture by other MGE routes can be explored.

Another key point in our study is the design and functionalization of CTCs capturing devices. The interactions between rare CTCs and solid surfaces are prone to interference by millions of WBCs as well as abundant biomolecules in blood samples. Therefore, surface functionalization of the CTCs capturing device with anti-fouling capability is indispensable. In lieu of the conventional CF-DBCO film, the nanoparticle-modified CF-NP-DBCO and CF-NP-SS-DBCO films are preferred in order to minimize non-specific absorption of WBCs. This is because the nanoparticles are composed of the zwitterionic sulfobetaine motif with good cytocompatibility and strong anti-fouling effects and promising for the modification of CTCs capturing devices. Furthermore, the CF-NP-SS-DBCO films are designed with breakable disulfide bonds, thereby facilitating the release of viable CTCs for various downstream applications including drug susceptibility test as demonstrated in our study.

### Data Availability

The authors declare that data supporting the findings of this study are available within the paper.

### Code Availability

The authors declare that code of this study is available within the paper.

## Methods

### Materials

Ethanol, methanol, dichloromethane, ethyl acetate, petroleum ether, pyridine, acetonitrile, *N,N*-dimethylformamide, triethylamine, D-mannosamine hydrochloride, cystamine dihydrochloride, trifluoroacetate, acetic anhydride, 4-dimethylaminopyridine (DMAP), SBMA, MAA, MBA, 2,2-azobisisobutyronitrile (AIBN), 1-ethyl-3-(3′-dimethylaminopropyl) carbodiimide (EDC), *N*-hydroxysuccinimide (NHS), DAPI, bovine serum albumin (BSA), and Histopaque-1077 solution were obtained from Sigma-Aldrich (USA). 2-NBDG, DBCO-NHS, DBCO-NH_2_, DBCO-biotin, and Alexa Fluor 488-labeled streptavidin were bought from Ruixi Co. Ltd. (China). Alexa Fluor 647-labeled CD45 probe, Alexa Fluor 488-labeled anti-pan-CK, Alexa Fluor 594-labeled anti-pan-CK were supplied by Univ-bio Co. Ltd. (China). Alexa Fluor 750-labeled CD45 gene probe, Alexa Fluor 594-labeled EpCAM gene probe, Alexa Fluor 594-labeled CK8, 18, 19 gene probes, Alexa Fluor 488-labeled vimentin gene probe, and membranes with a diameter of 8 μm (ISET standard) were bought from Xingyuan Co. Ltd. (China). Fluorouracil and oxaliplatin were provided by Shenzhen People’s Hospital.

### Chemical synthesis of Ac_4_ManNAc, Ac_4_ManNAz and DBCO-SS-NH_2_

The synthesis of Ac_4_ManNAc and Ac_4_ManNAz was according to a previously reported method.^98^ *Ac_4_ManNAc*: D-Mannosamine hydrochloride (215.63 mg, 1.00 mmol) was added to a flask containing pyridine (10 mL), acetic anhydride (1.02 g, 10.00 mmol), and DMAP (12.22 mg, 0.10 mmol). The mixture was stirred overnight, concentrated by rotary evaporation, and purified by column chromatography (ethyl acetate: petroleum ether = 1:1) to produce Ac_4_ManNAc (378.63 mg, 0.97 mmol, 97%) as a white solid.

#### Ac_4_ManNAz

A mixture of D-mannosamine hydrochloride (215.63 mg, 1.00 mmol), EDC (383.40 mg, 2.00 mmol), NHS (172.63 mg, 1.50 mmol) and triethylamine (0.5 mL) was stirred in methanol (10 mL). Azido acetic acid (121.27 mg, 1.20 mmol) was added and the mixture was stirred overnight, concentrated by rotary evaporation, and purified by column chromatography (methanol: dichloromethane = 5:1) to yield ManNAz (120.62 mg, 0.46 mmol, 46%) as a white solid. ManNAz (120.62 mg, 0.46 mmol) was added to a flask containing pyridine (5 mL), acetic anhydride (0.51 g, 5.00 mmol) and DMAP (6.11 mg, 0.05 mmol). The mixture was stirred overnight, concentrated by rotary evaporation, and purified by column chromatography (ethyl acetate: petroleum ether = 1:1) to produce Ac_4_ManNAz (189.36 mg, 0.44 mmol, 96%) as a white solid.

#### DBCO-SS-NH_2_

Di-tert-butyl decarbonate (2.18 g, 5.00 mmol) was added slowly to a *N,N*-dimethylformamide solution (20 mL) containing cystamine dihydrochloride (2.25 g, 10.00 mmol) and triethylamine (5 mL). The mixture was stirred overnight, concentrated by rotary evaporation, and purified by column chromatography (methanol: dichloromethane = 10:1) to produce cystamine mono-Boc ester (1.09 g, 4.31 mmol, 86%). A mixture of DBCO-NHS (402.40 mg, 1.00 mmol) and cystamine mono-Boc ester (252.40 mg, 1.00 mmol) was stirred in dichloromethane for 6 hours and purified by column chromatography (ethyl acetate: petroleum ether = 1:2) to produce DBCO-SS-NHBoc (539.72 mg, 0.92 mmol, 92%) as a white solid. DBCO-SS-NHBoc (539.72 mg, 0.92 mmol) was carefully treated by trifluoroacetate (1 mL) in dichloromethane (10 mL). DBCO-SS-NH_2_ (87.92 mg, 0.20 mmol, 22%) white solid was obtained after solvent evaporation, and then purified by column chromatography (methanol: dichloromethane = 4:1).

### MGE experiments

#### Cell culture

The human small cell lung cancer cell line (H524, suspended), human breast cancer cell line (MCF7), human T-mphoblastic cancer cell line (Jurkat, suspended), human cervical cancer cell line (HeLa), human non-small cell lung cancer cell line (A549), and human hepatoellular carcinomas cell lines (HepG2 and Huh7) were used in the cell experiments. All the cells were obtained from the Cell Bank of the Chinese Academy of Sciences. The H524 cells, MCF7 cells, A549 cells, HeLa cells and HepG2 cells were cultured by the Dulbecco’s modified eagle medium (DMEM) supplemented with 10% (v/v) fetal bovine serum (FBS) and 1% (w/v) penicillin/streptomycin. The Jurkat cells were cultured in the RPMI-1640 medium with 10% (v/v) FBS and 1% (w/v) penicillin/streptomycin. The Huh7 cells were cultured in DMEM with 10% (v/v) FBS and 1% (w/v) penicillin/streptomycin+NaHCO_3_. The PBMCs were cultured in the serum-free lymphocyte culture medium (Stemcell^®^).

#### MGE conditions in cell culture

Ac_4_ManNAz, Ac_4_ManNAc and DBCO-biotin were dissolved in DMSO at a concentration of 100 mM and stored at −20 °C. The solution was warmed to room temperature and added to cell culture medium with a certain concentration before use. The cancer cells were sub-cultured for at least 3 generations before use. The PBMCs were separated from fresh healthy blood by the Histopaque-1077 solution and SepMate^TM^-15 centrifuge tube according to manufacturer’s instruction.

#### MGE procedures

The cancer cells were seeded with a density of 0.3-0.4 million cells per mL in 4 mL of culture medium in a T25 flask. During cell culturing, Ac_4_ManNAz or Ac_4_ManNAc was added for a certain time to allow metabolic incorporation of *N-* azidoacetyl sialic acid to the glycoproteins of the cells. The PBMCs separated from 1 mL of fresh blood were seeded in 15 mL of the culture medium in a T25 flask and incubated with Ac_4_ManNAz or Ac_4_ManNAc under the same conditions.

#### Cell Viability

The cells were seeded with a density of 0.3 million cells per mL in 4 mL of the culture medium in a T25 flask with/without addition of Ac_4_ManNAz. After 12 hours, the number of cells was counted and the cell viability was calculated as the number of cells with Ac_4_ManNAz divided by the number of cells without Ac_4_ManNAz. Data were collected from 3 parallel tests.

#### Specific fluorescent staining of azido groups

Around 10000 cells with/without azido groups were washed with the labeling buffer (PBS solution with 2% FBS) and suspended in a tube or seeded onto a cell crawling sheet. The cells were incubated with DBCO-biotin (100 μM) in the labeling buffer for 1 hour at room temperature, washed 3 times with the labeling buffer (each time at least 10 minutes), and then incubated with Alexa Fluor 488-labeled streptavidin (50 μM) for 15 minutes in the dark. After fluorescent staining, the suspended cells were placed on a glass slide and the cells on the crawling sheet were used directly. All the cells were co-stained by DAPI for 3 minutes and the relative fluorescence intensity for each individual cell was monitored by automatic scanning fluorescence microscopy (Axio Imager Z2, Zeiss, Germany). Data points were collected in triplicate from 3 separate experiments.

*Note: In order to avoid false positive results from intracellular, the use of cell surface decontaminant reagents should be avoided*.

### Fabrication of multi-functional bio-orthogonal films

### CF

200 mg of chitosan (mw = 200,000) were dissolved in 10 mL of 90% acetic acid. 70 μL of the chitosan solution were added to round clean glass slides (d = 14 mm) and the film was prepared by spin-coating at 4000 revolutions per minute (rpm) for 15 seconds. The film was dried at 60 °C for 24 hours and 1 M NaOH was added to expose the active amino groups on the surface. The film was further dried at 60 °C for 24 hours to produce CF.

### CF-DBCO

CF was modified by 10 mM DBCO-NHS in ethanol overnight. The surface was washed with ethanol 3 times and PBS 3 times to produce CF-DBCO and then stored at 4 °C in darkness.

### Anti-fouling nanoparticles

The antifouling nanoparticles were prepared according to a modified protocol^92^. SBMA (1.00 g, 3.58 mmol), MAA (0.10 g, 1.16 mmol), MBA (0.10 g, 0.65 mmol), and AIBN (20.00 mg, 0.12 mmol) were added to acetonitrile (100 mL) and the mixture was heated to 110 °C for 30 minutes. After cooling to room temperature, the white nanoparticles were collected and washed with ethanol and water, centrifuged, and stored at 4 °C.

### CF-NP-DBCO

The nanoparticles were activated by 0.1 M EDC, 0.05 M NHS for 30 minutes and then added to CF. After 12 hours, the surface was washed with a mixture of ethanol/water 3 times and double-distilled water 3 times to remove uncombined nanoparticles and produce CF-NP-COOH (partly CF-NP-CO-NHS ester). The surface was re-activated by 0.1 M EDC and 0.05 M NHS for 30 minutes to make sure all the surface carboxyl groups were converted into NHS ester. Subsequently, DBCO-NH_2_ in DMSO (10 mM) was added slowly, shaken for 12 hours in the dark, and washed with ethanol/water 3 times and double-distilled water 3 times to produce CF-NP-DBCO. The CF-NP-DBCO films were stored at 4 °C in darkness.

### CF-NP-SS-DBCO

CF-NP-COOH was re-activated by 0.1 M EDC and 0.05 M NHS for 30 minutes to make sure all the surface carboxyl groups were converted into NHS ester. Afterwards, DBCO-SS-NH_2_ in DMSO (10 mM) was added slowly, shaken for 12 hours in darkness, and washed with ethanol/water 3 times and double-distilled water 3 times to produce CF-NP-SS-DBCO. The CF-NP-SS-DBCO films were stored at 4 °C in darkness.

### Characterization of bio-orthogonal films

#### Surface morphologies and chemical structures

The surface morphologies of the CF-DBCO, CF-NP-DBCO and CF-NP-SS-DBCO films were observed by SEM (SUPRA ^TM^ 55, Carl Zeiss, Germany) and the chemical structures were determined by XPS (ESCALab250Xi, Thermo Fisher Scientific, USA) and EDS (X-Max, Oxford Instruments, UK).

#### Capturing selectivity of azido-positive and azido-negative cancer cells

The cells were seeded with a density of 0.3∼0.4 million cells per mL in 4 mL of the culture medium. During cell culturing, Ac_4_ManNAz or Ac_4_ManNAc (100 μM) was added for 12 hours to allow metabolic incorporation of *N-*azidoacetyl sialic acid onto the glycoproteins of the cells. The treated cells were collected by centrifugation and washed by PBS for 3 times. Next, the bio-orthogonal films on a 24-well plate were added with 50000 cells in 1 mL of PBS and shaken at a speed of 20 rpm for 1 hour. The samples were rinsed with PBS at least 5 times and the captured cells were stained by DAPI and examined by fluorescent microscopy (BX51, Olympus Corporation, Japan) with 3 different views. The data were collected in triplicate from 3 separate experiments and the capturing rate was calculated as the number of captured cells divided by 50000.

#### Capturing selectivity of azido-positive cancer cells and PBMCs

The H524 cells were seeded with a density of 0.2∼0.3 million cells per mL in 4 mL of the culture medium and the PBMCs separated from 1 mL of fresh blood were seeded in 15 mL of the culture medium. During cell culturing, Ac_4_ManNAz (100 μM) was added for 12 hours to allow metabolic incorporation of *N-*azidoacetyl sialic acid onto the glycoproteins of the cells. The treated cancer cells and PBMCs were separately collected by centrifugation and washed with PBS 3 times. Afterwards, both kinds of cell suspensions (50000 H524 cells in 1 mL of PBS and 4∼8 million PBMCs in 1 mL of PBS) were prepared, added onto the bio-orthogonal films on a 24-well plate respectively, and then shaken at a speed of 20 rpm for 1 hour. The samples were rinsed with PBS at least 5 times and the captured cells were stained by DAPI and examined by fluorescent microscopy (BX51, Olympus Corporation, Japan) with 3 different views. The data were collected in triplicate from 3 separate experiments. The capturing rate was calculated as the number of captured cells divided by 50000 and that of PBMCs was calculated as the number of captured PBMCs divided by the number of PBMCs counted before capturing.

#### Release of the captured cancer cells

The cancer cells captured by the CF-NP-DBCO and CF-NP-SS-DBCO films underwent DAPI staining and examined by fluorescence microscopy. DTT (10 mM) was added to mediate disulfide reduction and release the captured cells. To ensure a sufficient reaction, the DTT solution was shaken at 80 rpm for 40 minutes and then the sample was washed gently with 2∼3 mL of PBS at least 5 times. The released cells were concentrated by centrifugation and re-suspended in 1 mL of PBS. 20 μL of the cell suspension were added to a glass slide and stained by DAPI to count the cell number. The released cells were further subjected to Calcein-AM/PI staining to evaluate the viability after release. The data were collected in triplicate from 3 separate experiments. The release rate was calculated as the number of released cells divided by the number of cells counted after capturing. The cell survival rate was calculated as the number of alive cells divided by the total number of released cells. For the evaluation of long-term cell viability, both the released cells and normal cells were added onto a 24-well plate at a density of 100000 per well and cultured for 1 and 3 days. At each time point, CCK-8 assay was performed according to manufacturer’s instruction. The long-term cell viability was calculated as the CCK-8 testing result at Day 3 divided by the CCK-8 testing result at Day 1.

#### Comparison of capturing capability between ISET and bio-orthogonal films

10000 HeLa or H524 cells were dispersed in 10 mL of PBS, and then purified by the ISET standard membrane according to manufacturer’s instruction. As for the bio-orthogonal capture, 10000 HeLa or H524 cells labeled by Ac_4_ManNAz were dispersed in 1 mL of PBS, added onto the bio-orthogonal films on a 24-well plate, and then shaken at a speed of 20 rpm for 1 hour, followed by PBS washing at least 5 times. The data were collected in triplicate and the capturing rate was calculated as the number of captured cells divided by 10000.

#### Capture of artificial CTCs in blood samples

10000 HeLa or H524 cells were dispersed in 1 mL of PBS, and 10 μL of the cell suspensions were added into 1 mL of fresh blood separately to obtain artificial CTCs. Each sample was pretreated with the ACK lysis buffer at 0 °C for 10 minutes and collected by centrifugation under 300 g. After washing with PBS containing 2% FBS at 120 g once, the remaining cells were cultured in the serum-free lymphocyte culture medium containing 100 μM Ac_4_ManNAz for 12 hours. The treated cells were washed with PBS 3 times, added to CF-NP-DBCO on a 24-well plate, and shaken at 20 rpm for 1 hour. The samples were rinsed with PBS at least 5 times and the cells were fixed by 4% paraformaldehyde and Triton X-100 (0.1% in H_2_O, 5 minutes). Finally, the fixed cells were treated by Alexa Fluor 594-labeled pan-CK probe (50 μM, 1 hour), Alexa Fluor 750-labeled CD45 probe (50 μM, 3 hours) and DAPI (10 μg/mL, 3 minutes), respectively. An automatic scanning fluorescent microscope (Axio Imager Z2, Zeiss, Germany) was used to identify the captured cancer cells as the Nucleus+/CD45-/pan-CK+. The data were collected 9 times for each cell lines and the capturing rate was calculated as the number of captured cancer cells divided by 100.

### Clinical CTCs detection

#### Study approval

This clinical study was approved by the Ethics Committee of Shenzhen People’s Hospital, China (approval number: KY-LL--2020157-02). All the participants provided written informed consent for sample collection and subsequent analysis.

#### Cells isolated by ISET and identified by nucleus/CD45/clinical-marker/azido staining

The ethylenediaminetetraacetic acid (EDTA) anticoagulated whole blood samples were obtained from different cancer patients in Shenzhen People’s Hospital. Each blood sample was pretreated with ACK lysis buffer at 0 °C for 10 minutes and purified by the ISET standard membrane according to manufacturer’s instruction. The cells remaining on ISET membrane were cultured in the serum-free lymphocyte culture medium containing 100μM Ac_4_ManNAz for 12 hours. Afterwards, the membrane was washed with PBS 5 times and the cells were fixed by 4% paraformaldehyde. Finally, the fixed cells were treated by DBCO-biotin (100 μM, 1 hour), Alexa Fluor 488-labeled streptavidin (50 μM, 15 minutes), Triton X-100 (0.1% in H_2_O, 5 minutes), Alexa Fluor 594-labeled pan-CK probe (50 μM, 1 hour) or Alexa Fluor 594-labeled EpCAM-CKs probe (a mixture of EpCAM, CK8, CK18, CK19 probe, 50 μM for each, 1 hour), Alexa Fluor 750-labeled CD45 probe (50 μM, 3 hours) and DAPI (10 μg/mL, 3 minutes), respectively. An automatic scanning fluorescent microscope (Axio Imager Z2, Zeiss, Germany) was used to identify CTCs as Nucleus+/CD45-/Clinical marker+.

#### Cells captured by the bio-orthogonal films and identified by Nucleus/CD45/EpCAM/vimentin staining

The EDTA anticoagulated whole blood samples were obtained from different cancer patients in Shenzhen People’s Hospital and healthy people. Each blood sample was pretreated with the ACK lysis buffer at 0 °C for 10 minutes and collected by centrifugation under 300 g. After washing with PBS containing 2% FBS at 120 g once, the remaining cells were cultured in the serum-free lymphocyte culture medium containing 100 μM Ac_4_ManNAz for 12 hours. The treated cells were washed with PBS 3 times, added to CF-NP-DBCO on a 24-well plate, and shaken at 20 rpm for 1 hour. The samples were rinsed with PBS at least 5 times and the cells were fixed by 4% paraformaldehyde and Triton X-100 (0.1% in H_2_O, 5 minutes). Finally, the fixed cells were treated with Cy3-labeled EpCAM probe (50 μM, 1 hour), Alexa Fluor 488-labeled vimentin probe (50 μM, 1 hour), Alexa Fluor 750-labeled CD45 probe (50 μM, 3 hours) and DAPI (10 μg/mL, 3 minutes) according to manufacturer’s instruction. An automatic scanning fluorescent microscope (Axio Imager Z2, Zeiss, Germany) was used to identify the captured CTCs as Nucleus+/CD45-/EpCAM+ and/or vimentin+.

#### Direct comparison of CTCs detection between our strategy and TumorFisher^®^ detection system

The EDTA anticoagulated whole blood samples were obtained from different cancer patients in Shenzhen People’s Hospital and healthy volunteers. Each blood sample was halved and parallelly processed by TumorFisher® and our detection workflow using CF-NP-DBCO films as described above. Afterward, all the CTCs captured by different systems were examined by immunohistochemical staining.

#### Release of CTCs and glycolytic activity detection

The EDTA anticoagulated whole blood samples were obtained from different patients with stage III colon cancers in Shenzhen People’s Hospital. Each blood sample was processed by our detection workflow using CF-NP-SS-DBCO films as described above. Afterward, the glucose-free DMEM medium containing DTT (10 mM) was added for 40 minutes to allow the release of captured CTCs. The released cells were collected and cultured with the glucose-free DMEM medium containing 2-NBDG (100 μM), Alexa Fluor 647-labeled CD45 probe (10 μM), and Hoechst-33342 (10 μM) for 15 minutes to identify CTCs as Nucleus+/CD45- with the glycolytic activity being examined by 2-NBDG. After fluorescent staining, the medium (20 μL) was added to the confocal dish with a 3 mm x 3 mm hole and observed under a confocal microscope (STEDYCON, Leica, Germany).

#### Drug susceptibility test using released CTCs

The EDTA anticoagulated whole blood sample was obtained from a patient with stage III colon cancer in Shenzhen People’s Hospital. The blood sample was evenly divided into 3 potions, and processed by our detection workflow using CF-NP-SS-DBCO films as described above. In the test of drug susceptibility, fluorouracil (20 mM) or oxaliplatin (20 mM) was added during the MGE process. Afterward, the glucose-free DMEM medium containing DTT (10 mM) was added to each group for 40 minutes to allow the release of captured CTCs. The released CTCs in different groups were identified and examined for the glycolytic activity as described above. ImageJ software was used to analyze the fluorescence intensity of CTCs after different treatments, and the relative fluorescence intensity (fluorescence intensity/cell area) of each 2-NBDG stained CTC was calculated.

### Statistical analysis

The cell experiments were performed at least in triplicate with the results being presented as mean + standard deviation. The student’s *t-test* was performed to determine the levels of statistical significance among different groups. A difference of \**p* < 0.05 was considered to be significant and that of \*\**p* < 0.01 or \*\*\**p* < 0.001 was considered to be highly significant.

## Acknowledgements

This work was financially supported by the National Key Research and Development Program of China (2021YFB3800800), National Natural Science Foundation of China (81903057, 82073284, 32000962, 82074133 and 82272157), Shenzhen Science and Technology Research Funding (JCYJ20200109115601720 and JSGG20200225152648408), Guangdong Basic and Applied Basic Research Foundation (2021A1515012163), China Postdoctoral Science Foundation (2020M683200), Hong Kong PDFS - RGC Postdoctoral Fellowship Scheme (PDFS2122-1S08 and CityU 9061014), as well as Hong Kong HMRF (Health and Medical Research Fund) (2120972 and CityU 9211320). We also acknowledge helpful discussions with Dr. Ya Zhao, Dr. Guofen Song, Dr. Ang Gao and Dr. Shi Mo. Moreover, we acknowledge the helpful technique support from Dr. Xiaoyu Pu from Xingyuan Co. Ltd. (China).

## Author contributions

Z. L. performed the chemical synthesis and material preparation experiment, Z. L., X. R. and L. X. performed the cell experiments, Y. Z. collected clinical blood samples. Z. L. and D. Z. performed the clinical tests, Z. L., X. R., Z. J., P. L. and L. T. analyzed the data, W. L., Y. C. and H. W. contributed to the theoretical interpretation of the data, Z. L., X. R., W. L., P. K. C. and H. W. wrote the manuscript with comments from all co-authors. Z. L., W. L., Y. C., and H. W. conceived and supervised the project.

