## Supplemental Information for "Beyond enumeration: Phenotype independent “labeling-capture-release” process enabling precise detection of circulating tumour cells and downstream applications"

*#These authors contributed equally*

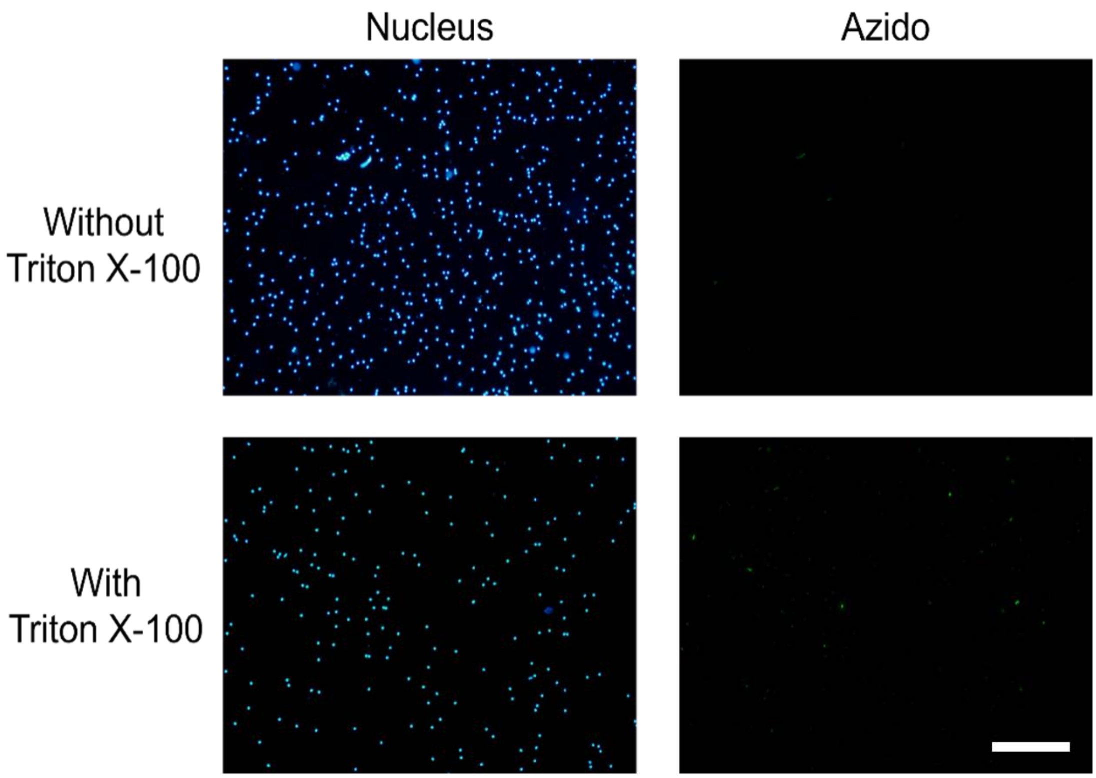

**Supplementary Figure 1.** Fluorescent images of MGE-labeled PBMCs with/without the treatment of surface decontaminant reagent (scale bar = 200 μm). Some PBMCs can absorb Ac4ManNAz, but cannot deliver the azido groups to the cell surface. In order to specifically label the azido groups on cells, labeling has to be performed without the surface decontaminant reagent.

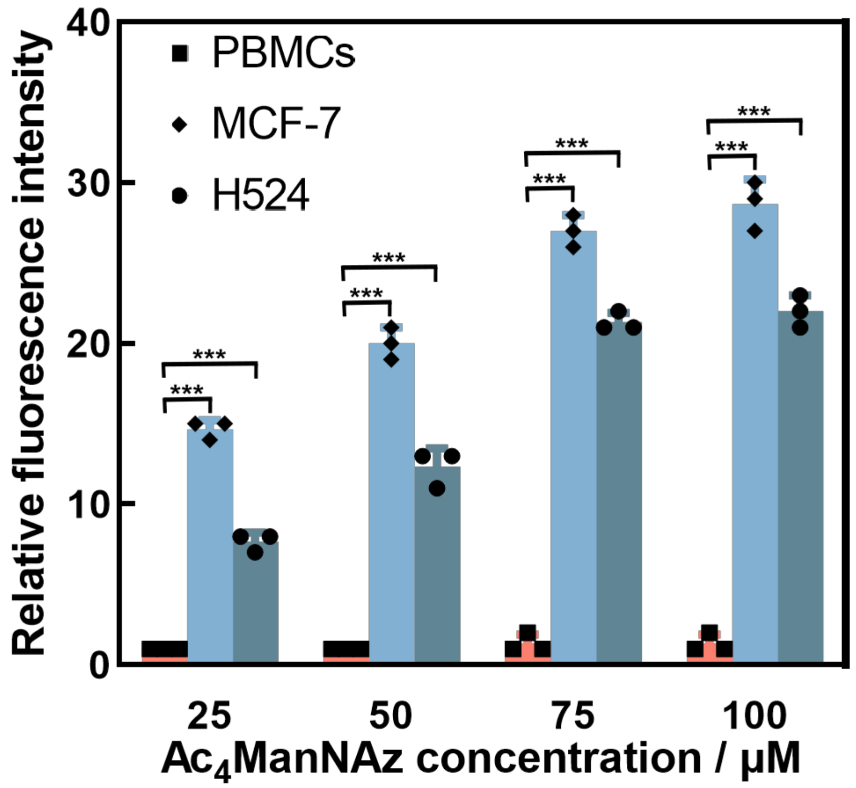

**Supplementary Figure 2.** Sugar concentration-cell surface fluorescence intensity histogram (*** denotes $p < 0.001$).

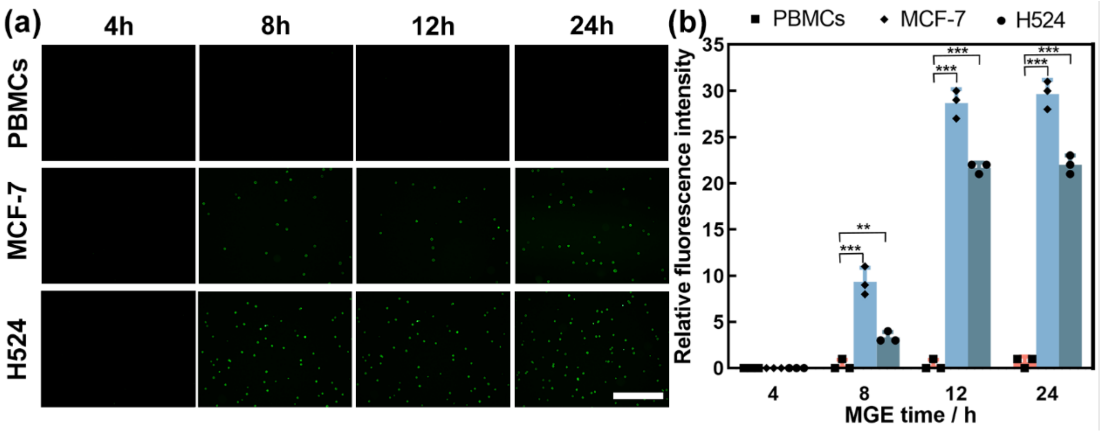

**Supplementary Figure 3.** (a) Fluorescent images of different cells for different MGE time (scale bar = 200 μm); (b) MGE time-cell surface fluorescence intensity histogram (** denotes $p < 0.01$ and *** denotes $p < 0.001$).

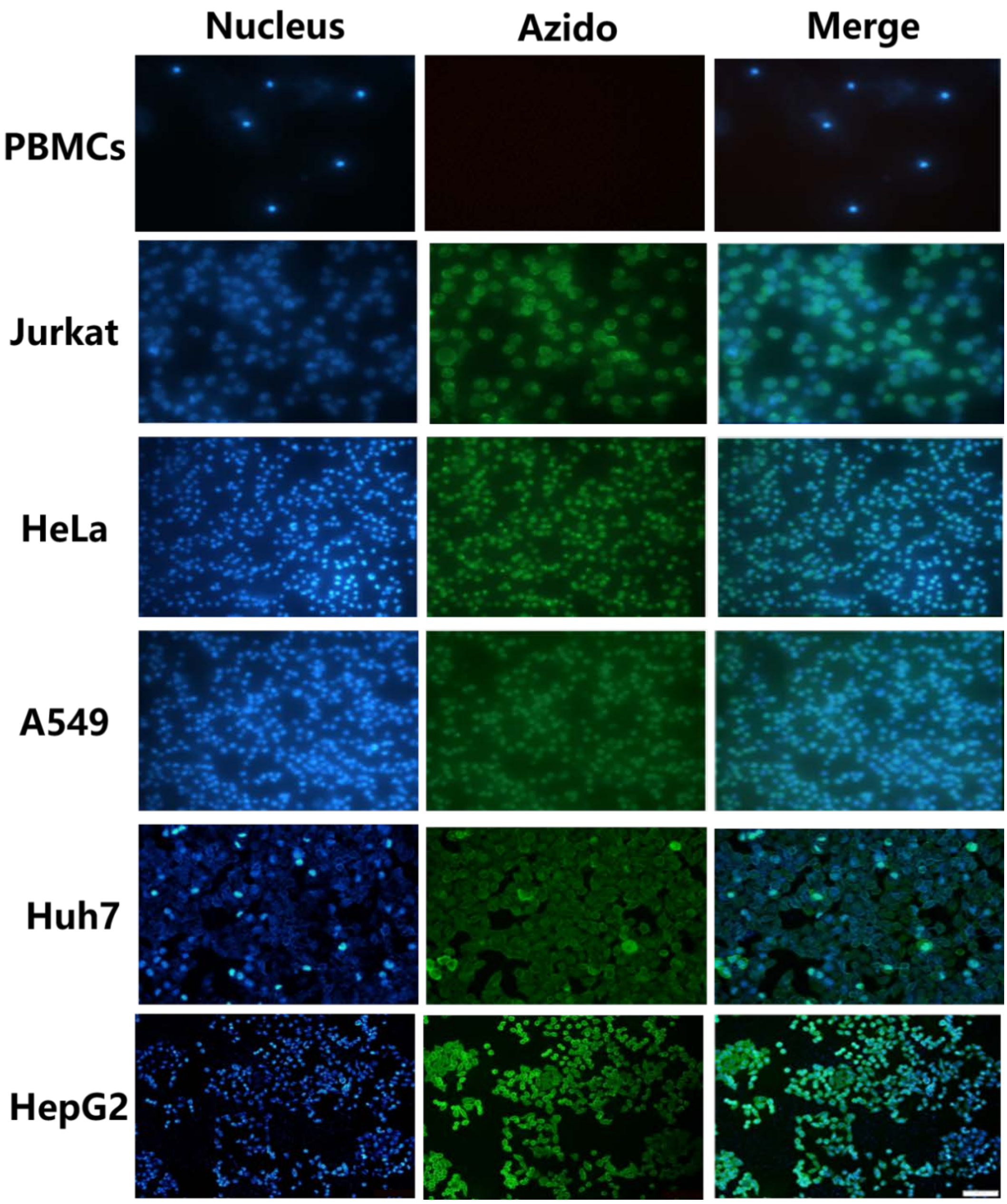

**Supplementary Figure 4.** Fluorescent images of MGE-labeled cancer cells and PBMCs (scale bar = 200 μm) showing that the cancer cells are selectively modified by MGE while normal blood cells are not.

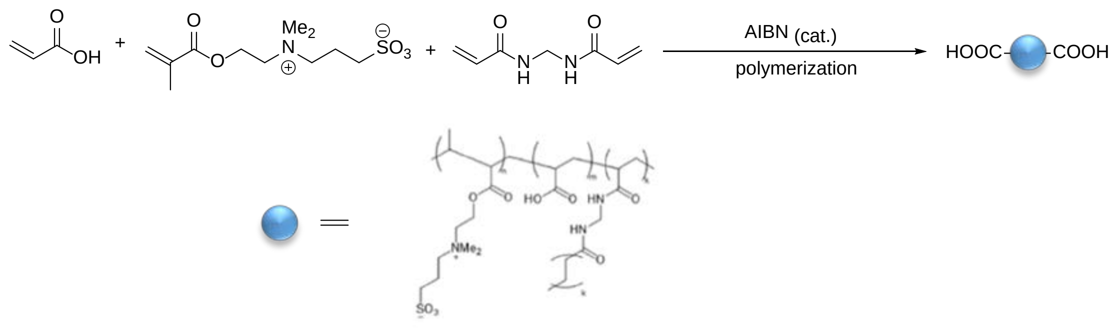

**Supplementary Figure 5.** Synthesis of anti-fouling nanoparticles.

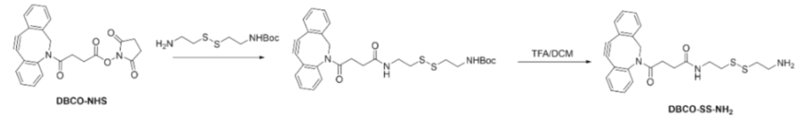

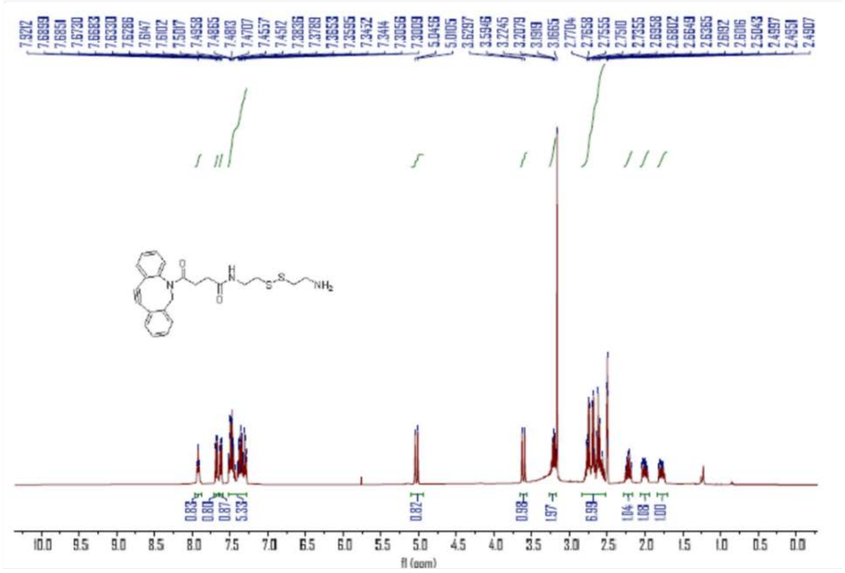

$^1$H-NMR Spectrum of DBCO-SS-NH$_2$

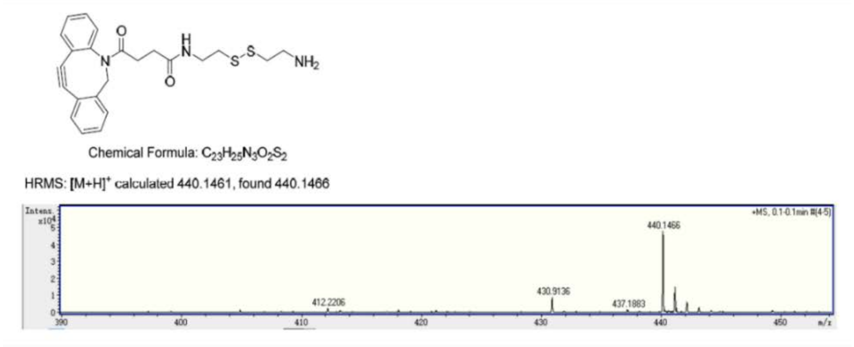

HRMS Spectrum of DBCO-SS-NH$_2$

**Supplementary Figure 6**. Synthesis and characterization of DBCO-SS-NH$_2$: $^1$H NMR (400 MHz, dimethylsulfoxide-d6) δ 7.91 (1H, t), 7.71-7.25 (8H, m), 5.02 (1H, d), 3.61 (1H, d), 3.16 (2H, m), 2.77-2.52 (7H, m), 2.24 (1H, dd), 2.01 (1H, dd), 1.31 (1H, m); High-resolution mass spectroscopy (HRMS), $[M+H]^+$, calculated 440.1461, found 440.1466.

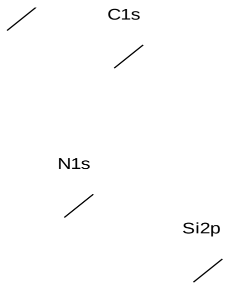

| | |
|---|---|
| O | 22.48 |
| C | 58.57 |
| N | 7.53 |
| Si | 11.43 |
| S | 0 |

**CF-DBCO**

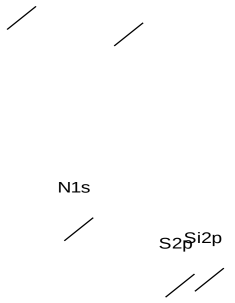

| | |
|---|---|
| O | 22.8 |
| C | 62.38 |
| N | 6.02 |
| Si | 6.73 |
| S | 2.07 |

**CF-NP-DBCO**

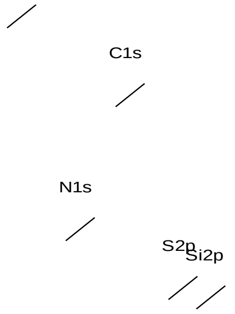

| | |
|---|---|
| O | 25.76 |
| C | 62.37 |
| N | 5.83 |
| S | 2.69 |
| Si | 3.35 |

**CF-NP-SS-DBCO**

**Supplementary Figure 7**. XPS analysis of different bio-orthogonal films.

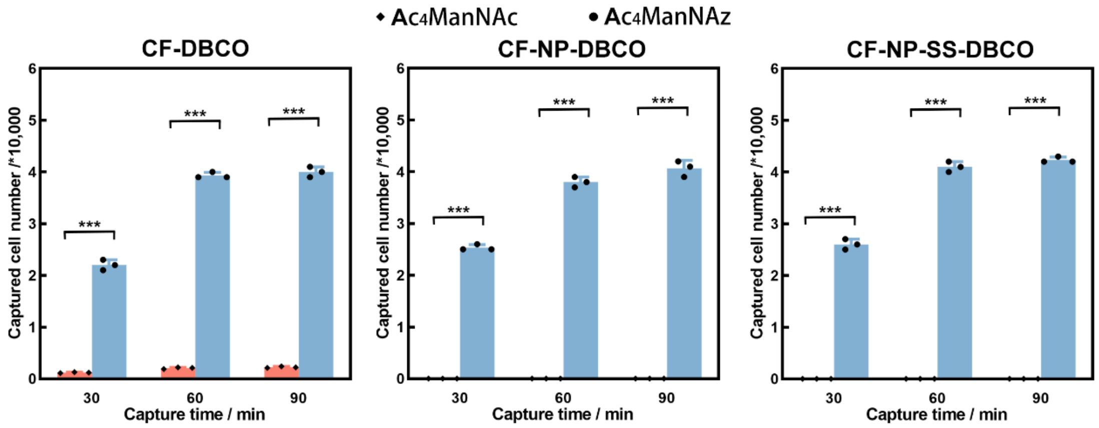

**Supplementary Figure 8**. Capture time-captured cell number (from a total of 50000 cells) of MGE-treated H524 cells with/without azido groups on the bio-orthogonal films (*** denotes $p < 0.001$).

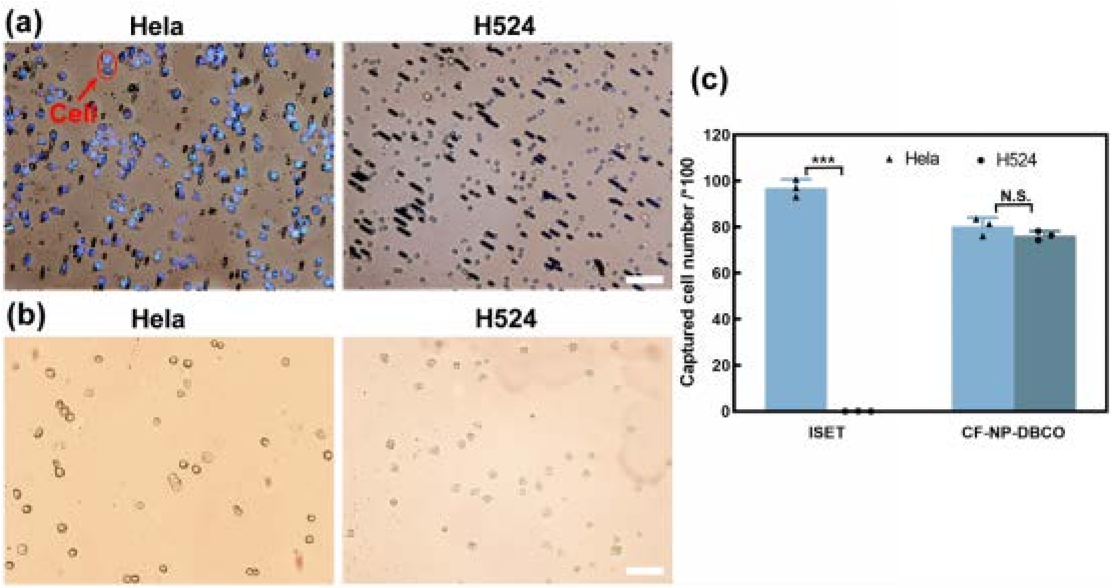

**Supplementary Figure 9**. Comparison of capturing capability between the bio-orthogonal films and commercial ISET. (a) HeLa cells (left) and H524 cells (right) captured by ISET standard membrane (scale bar = 50 μm); (b) HeLa cells (left) and H524 cells (right) captured by bio-orthogonal films (scale bar = 50 μm); (c) Statistical analysis (*** denotes $p < 0.001$).

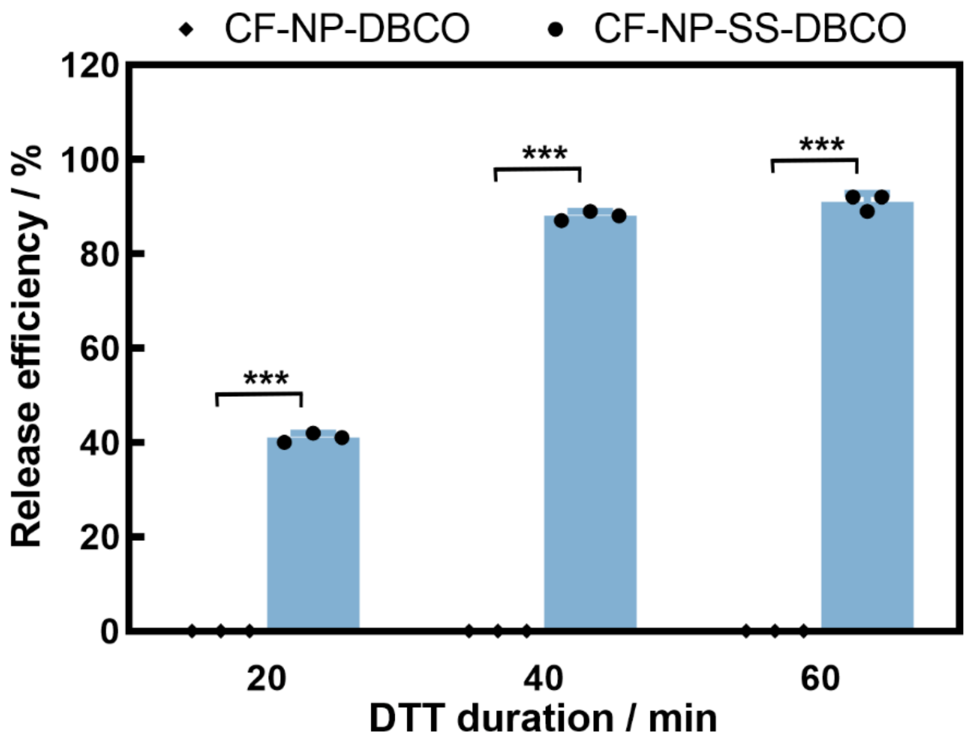

**Supplementary Figure 10**. DTT treatment duration-H524 cells release efficiency of CF-NP-DBCO and CF-NP-SS-DBCO (*** denotes $p < 0.001$).

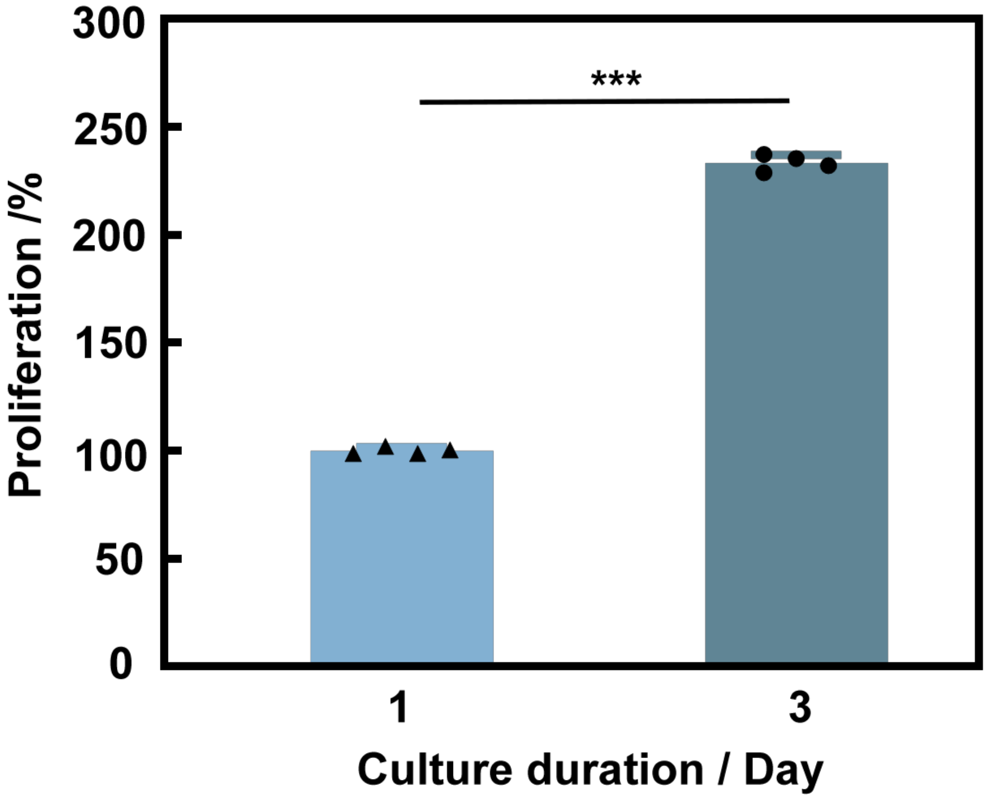

**Supplementary Figure 11**. Long-term viability of H524 cells by normal cultivation (*** denotes $p < 0.001$).

(a) H524 artificial CTC sample

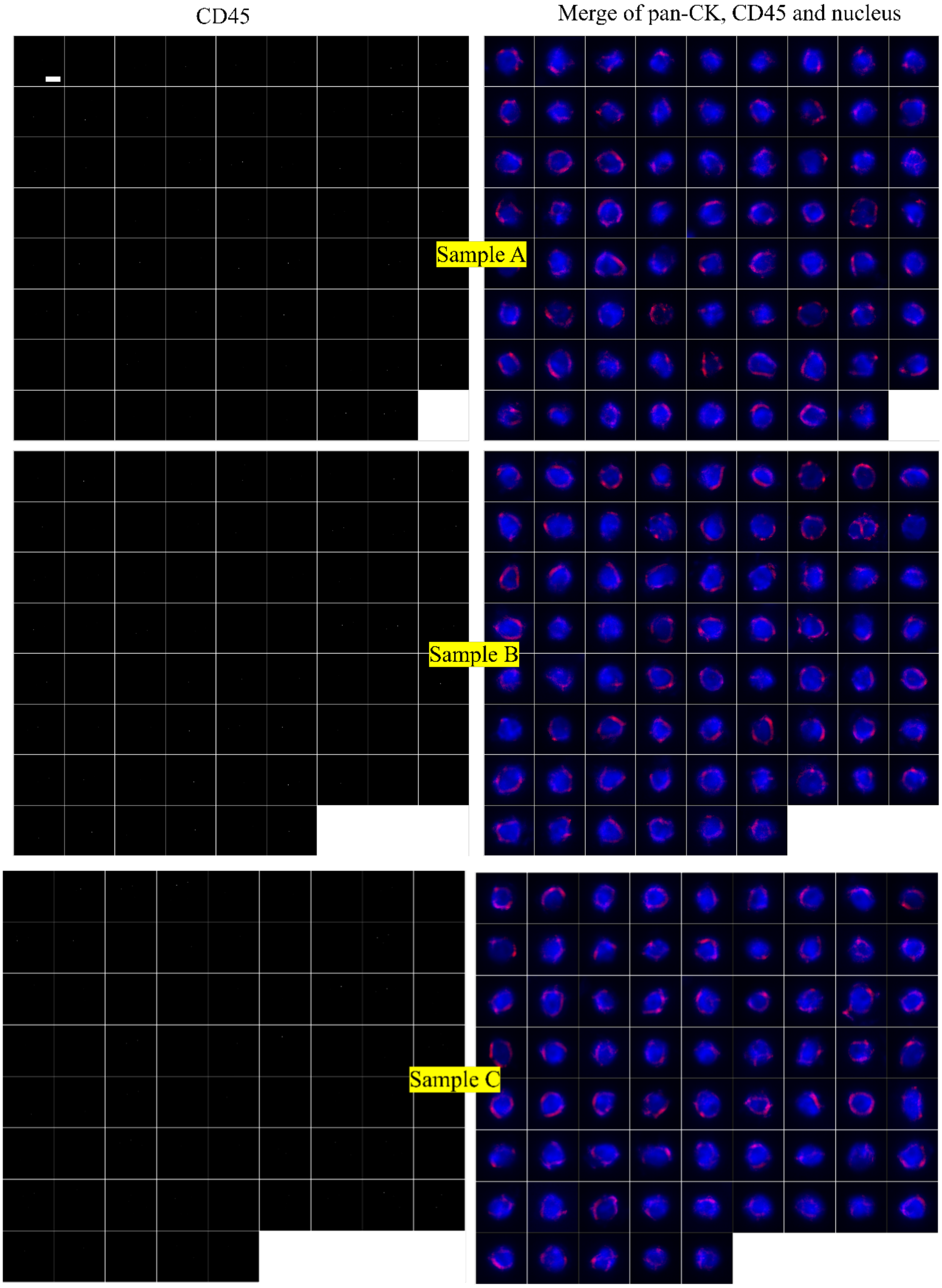

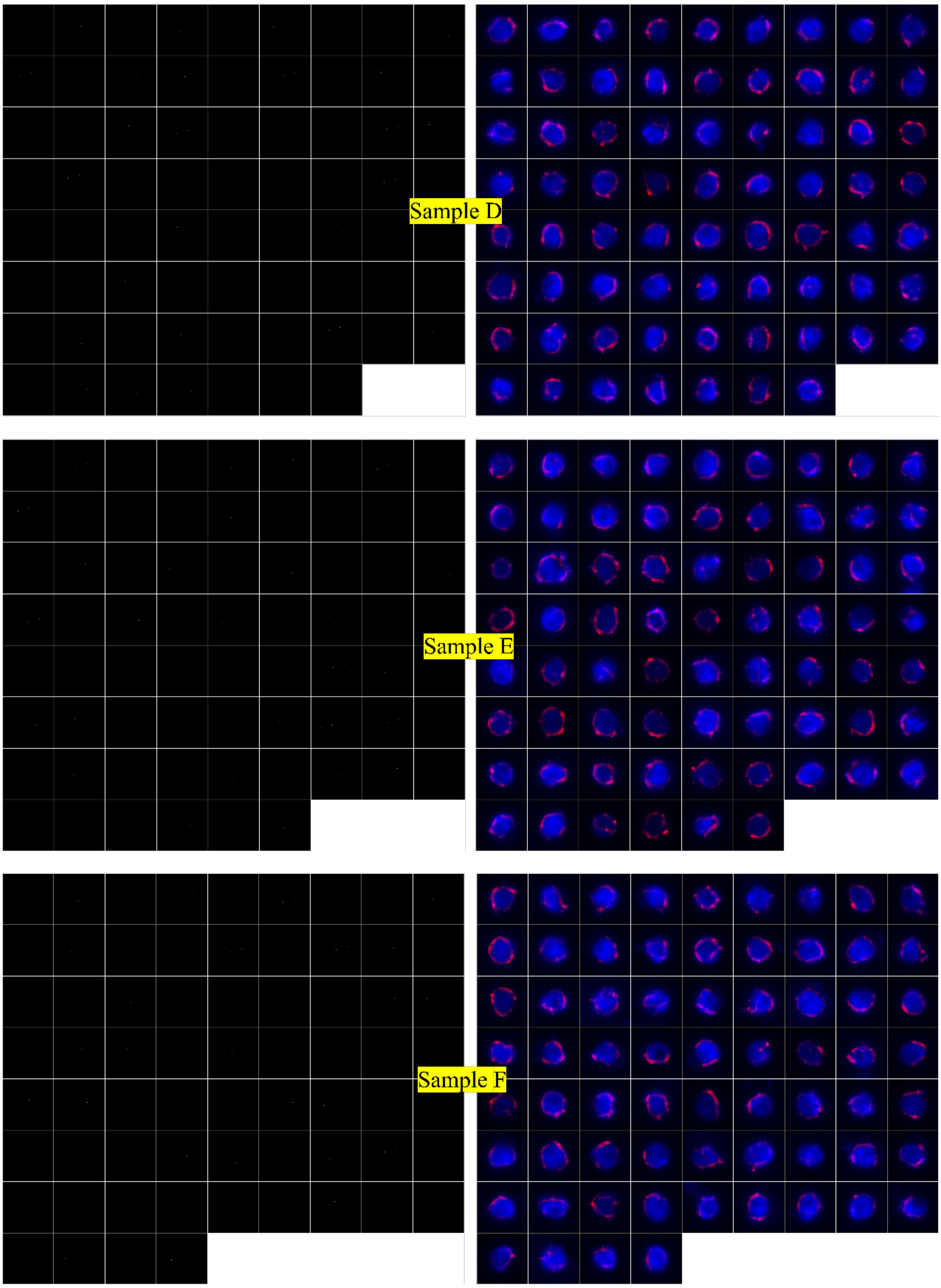
Sample D
Sample E
Sample F

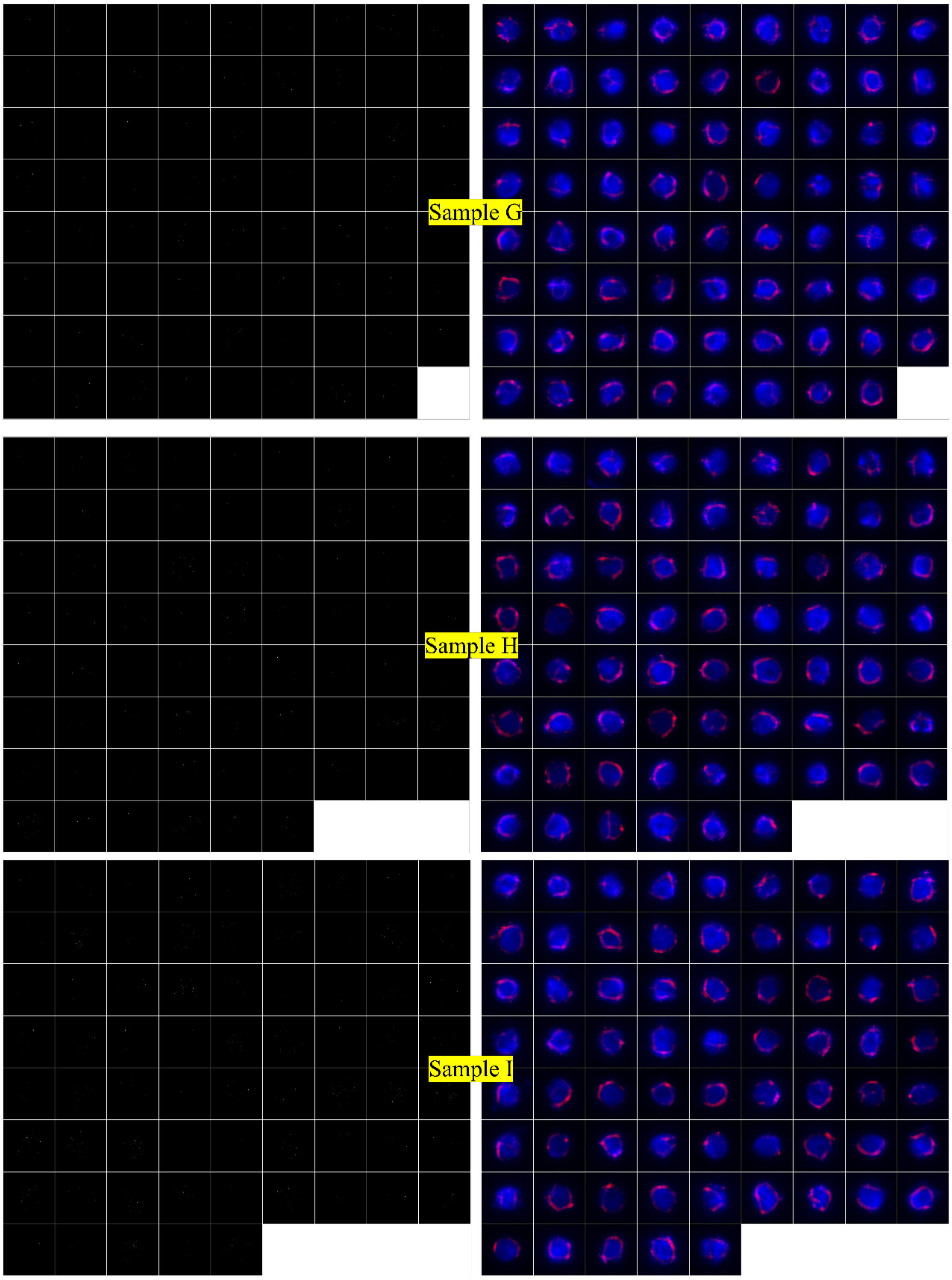
Sample G
Sample H
Sample I

(b) HeLa artificial CTC sample

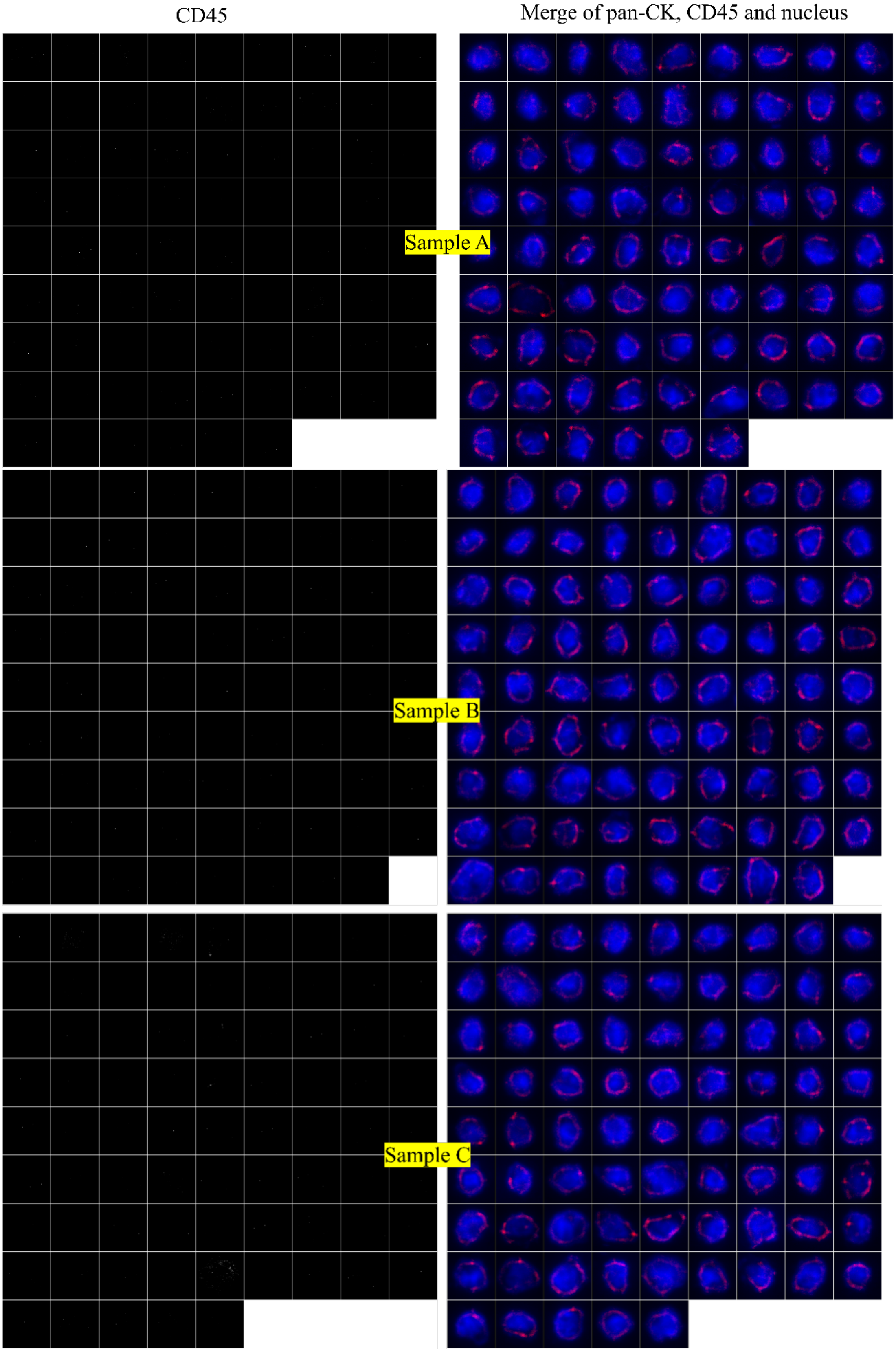

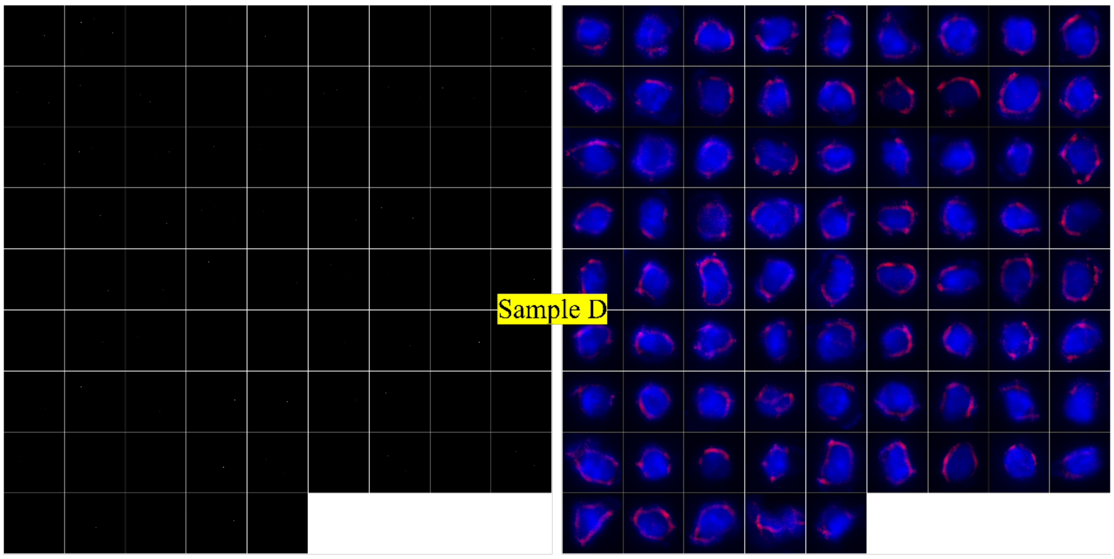
Sample D

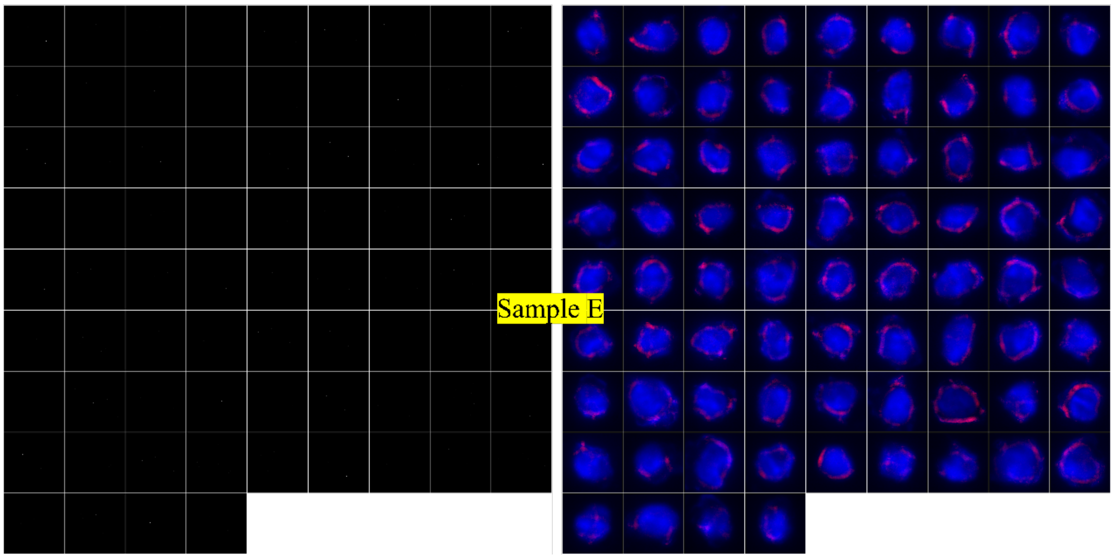
Sample E

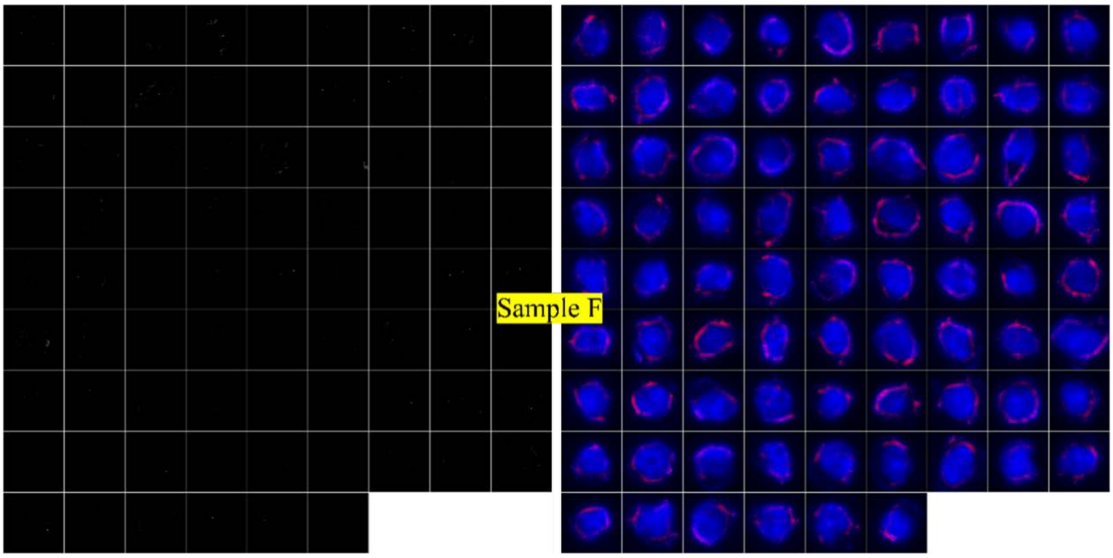
Sample F

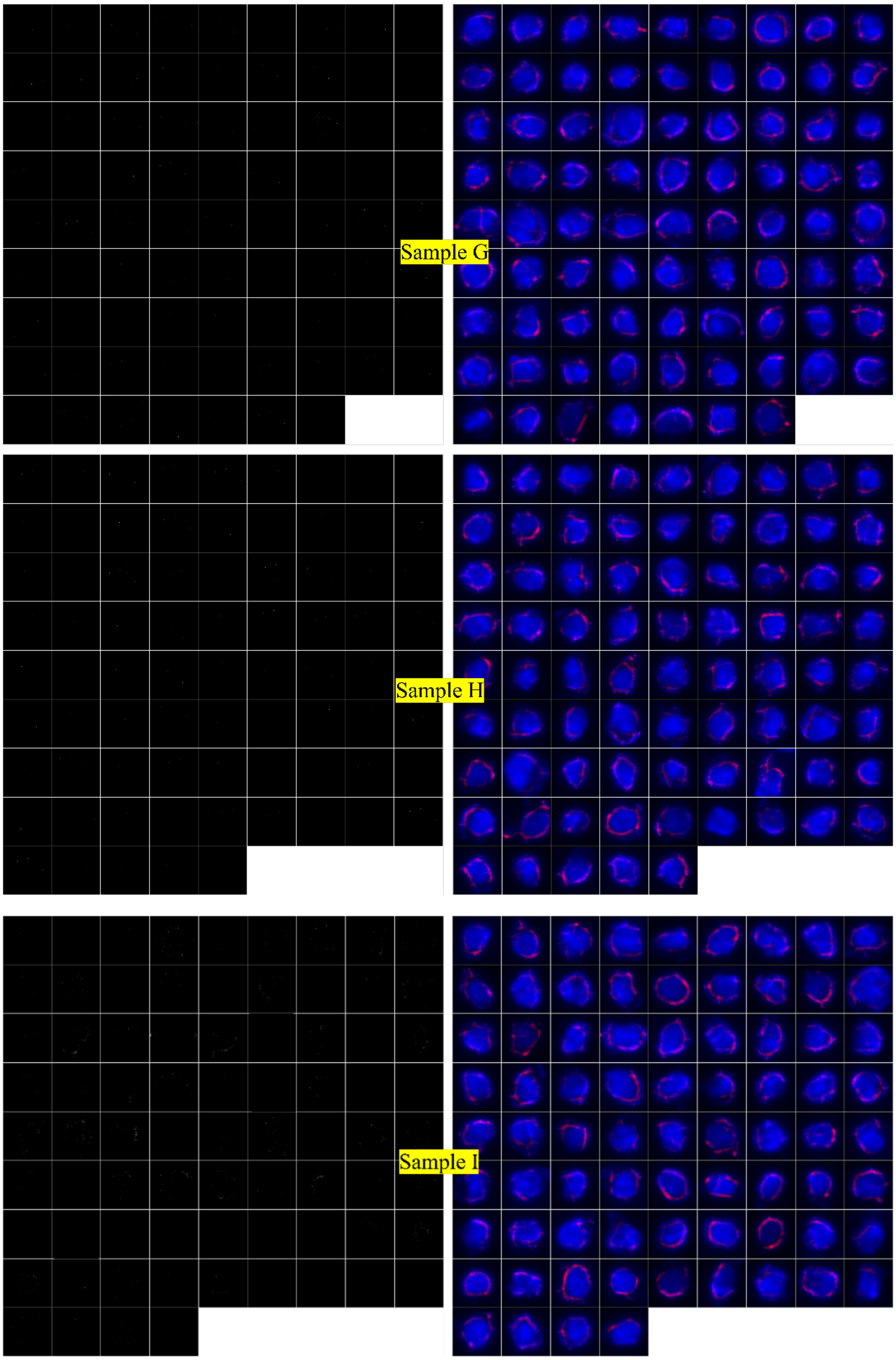
Sample G
Sample H
Sample I

**Supplementary Figure 12**. Fluorescent pictures of artificial CTCs (100 cancer cells spiked in 1 mL of blood samples) captured by the bio-orthogonal films and identified by the nucleus (blue), CD45 (white), and pan-CK (red) staining (scale bar = 5 μm). (a) Captured H524 cells in blood samples and (b) Captured HeLa cells in blood samples.

***Patient information: male, 73 years old, liver cancer.***

**(a) Picture by 590 nm and 490 nm excitation luminescence**

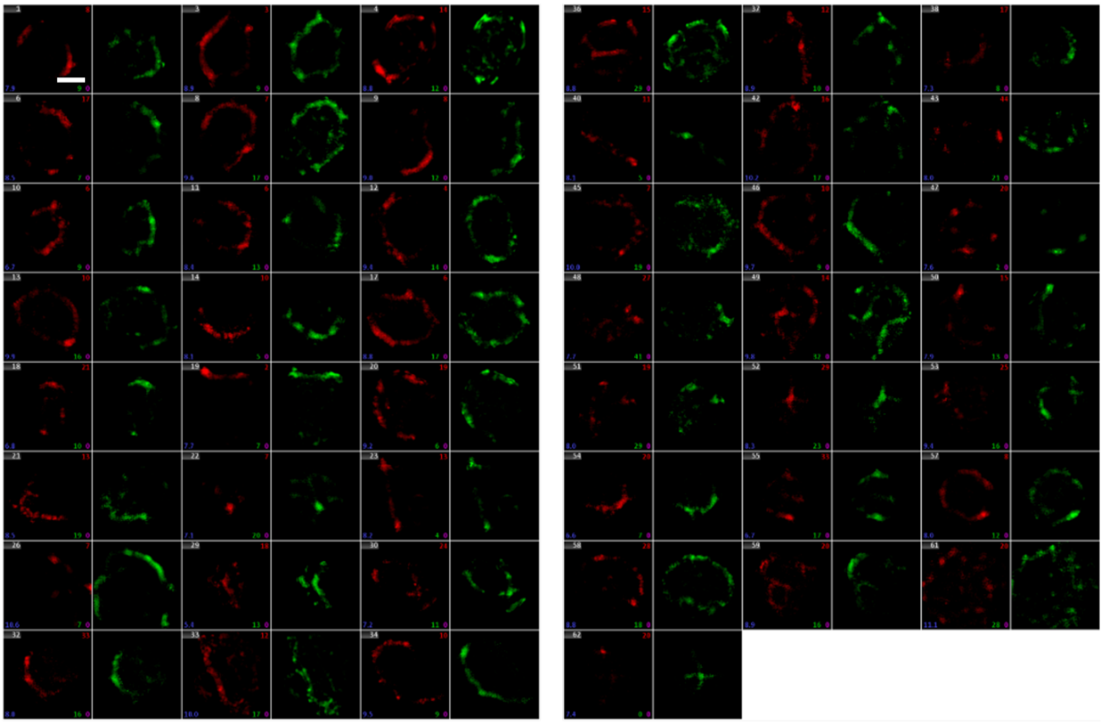

**(b) Picture by750 nm excitation luminescence**

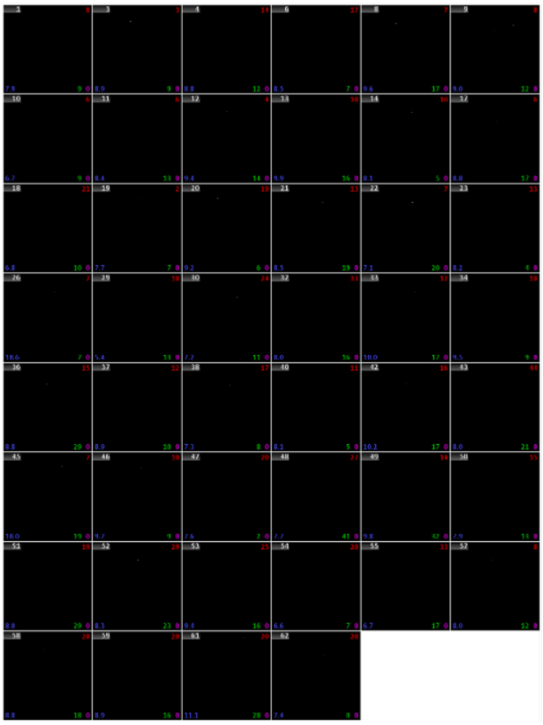

**(c) Merge**

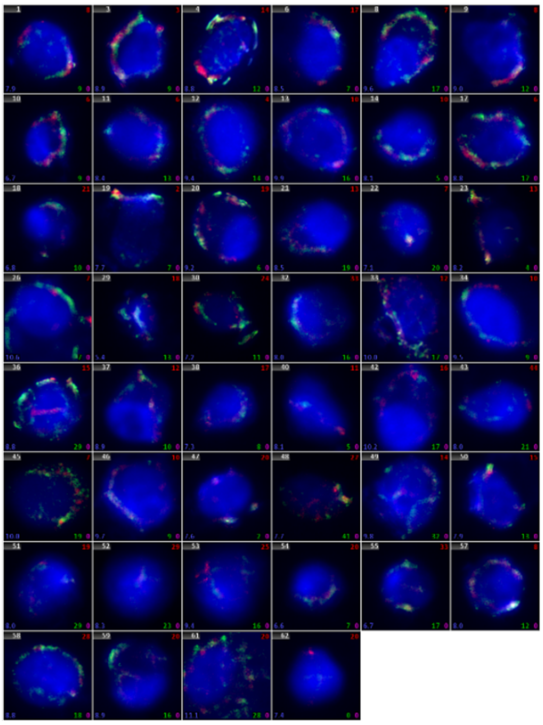

**Supplementary Figure 13**. Fluorescent pictures of ISET-separated clinical CTCs modified by MGE with pan-CK as the positive CTCs biomarker (scale bar = 5 μm). (a) Red referring to pan-CK and green color representing the azido group; (b) White color refers to CD45; (c) Merged images of pan-CK, azido group, CD45, and nucleus.

*Patient information: Sample A, Male, 52 years old, liver cancer;*
*Sample B, Female, 61 years old, lung cancer; Sample C, Male, 45 years old, nasopharyngeal cancer.*

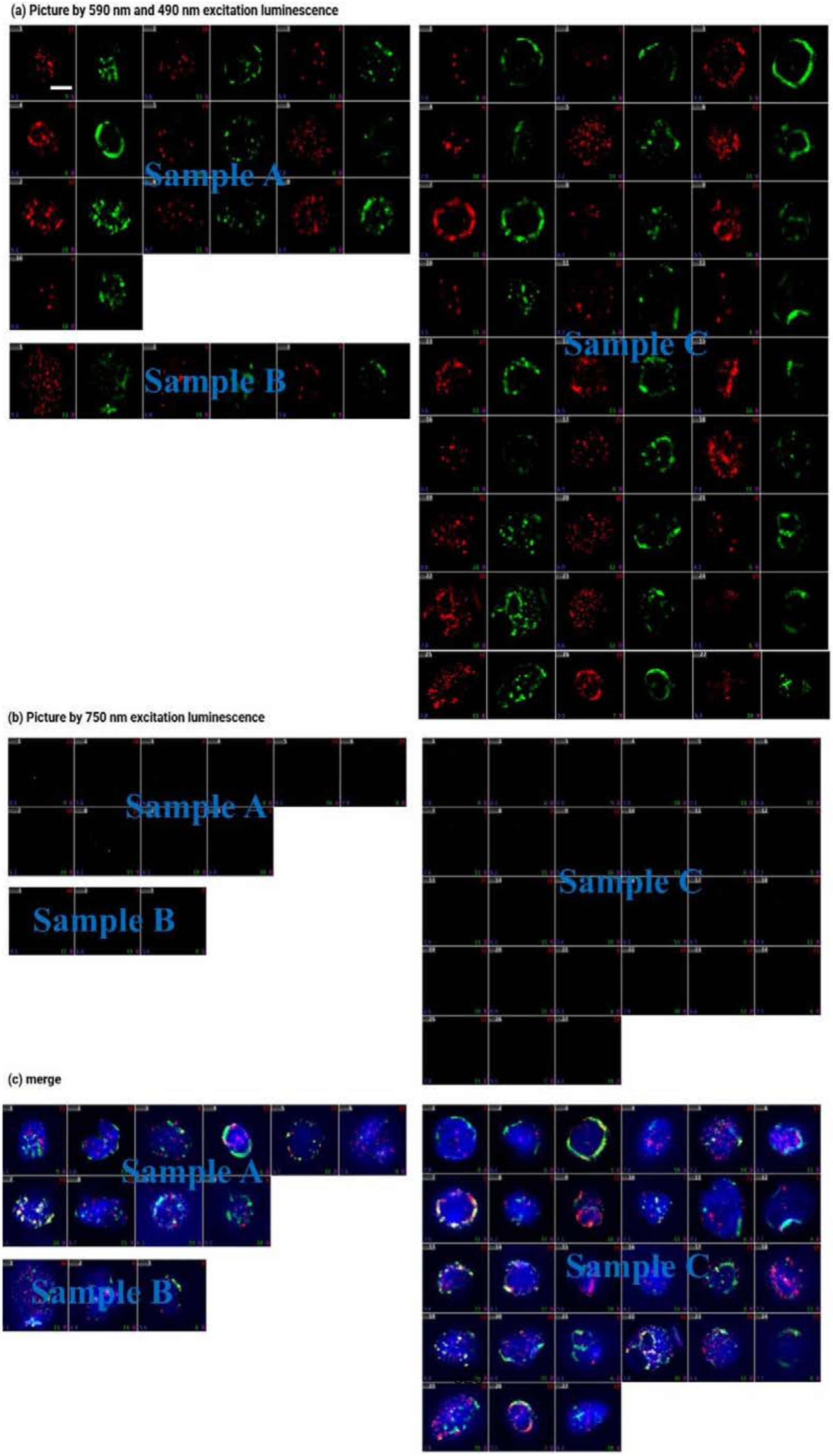

**Supplementary Figure 14**. Fluorescent pictures or ISET-separated clinical CTCs modified by MGE with EpCAM-CKs as the positive CTCs biomarker (scale bar = 5 μm). (a) Red referring to EpCAM-CKs and green referring to the azido group; (b) White color refers to CD45; (c) Merged images of EpCAM-CKs, azido group, CD45 and nucleus.

**P1**

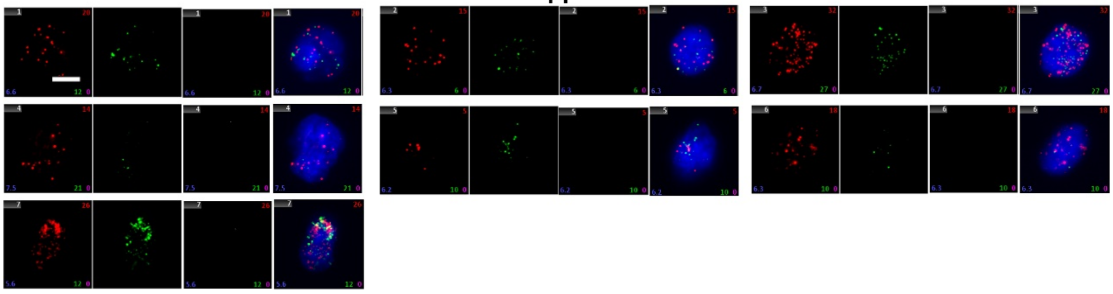

**P2**

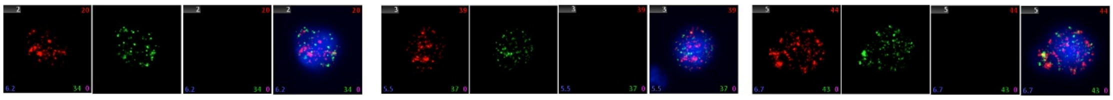

**P3**

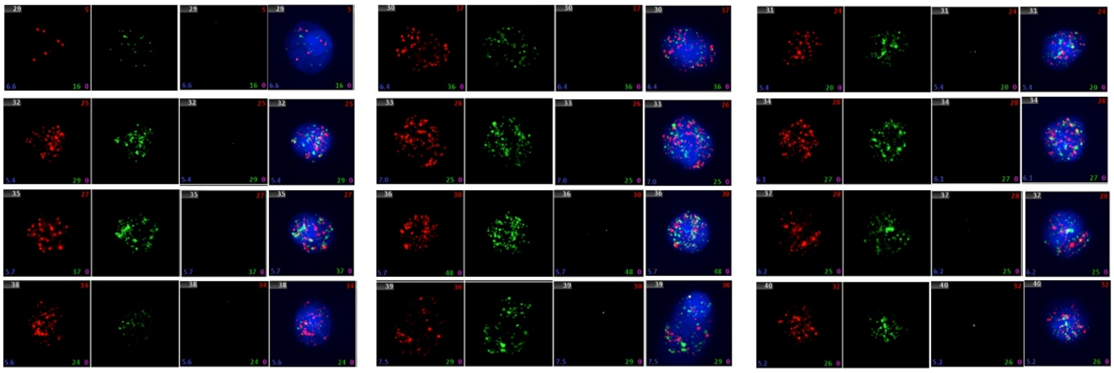

**P4**

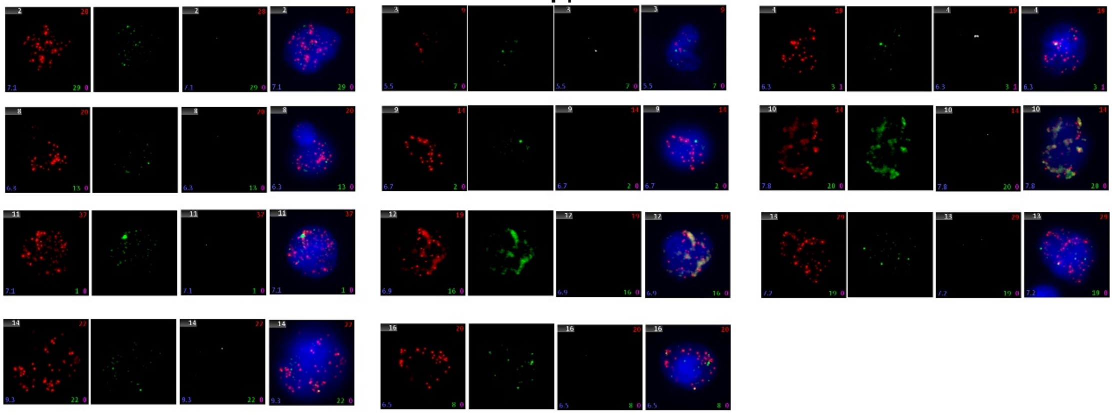

**P5**

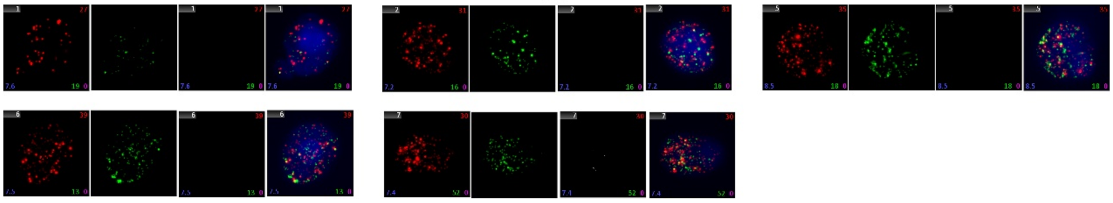

P6

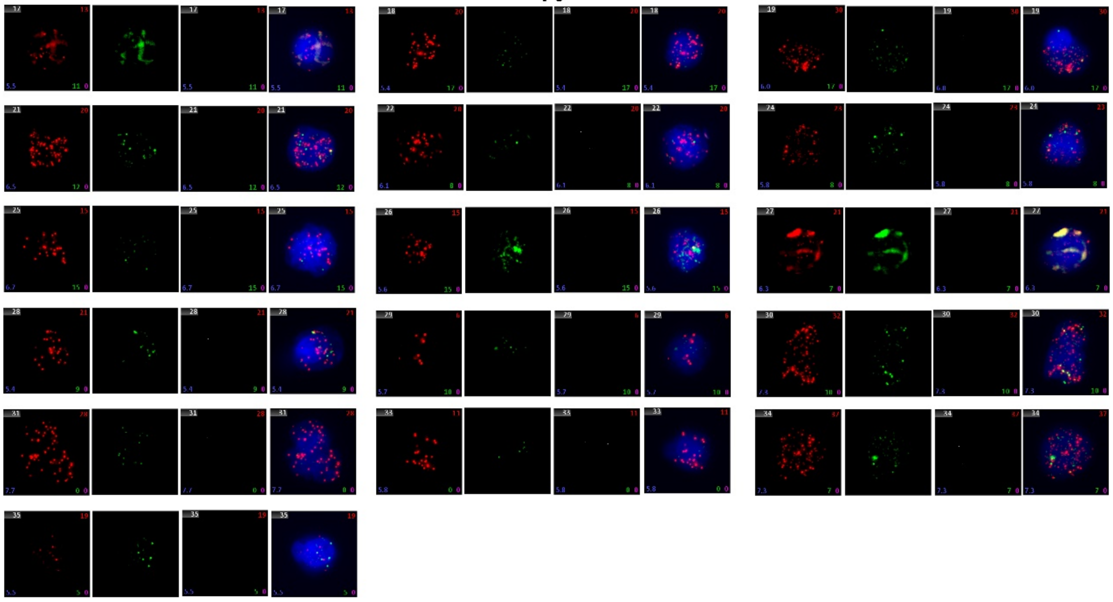

P7

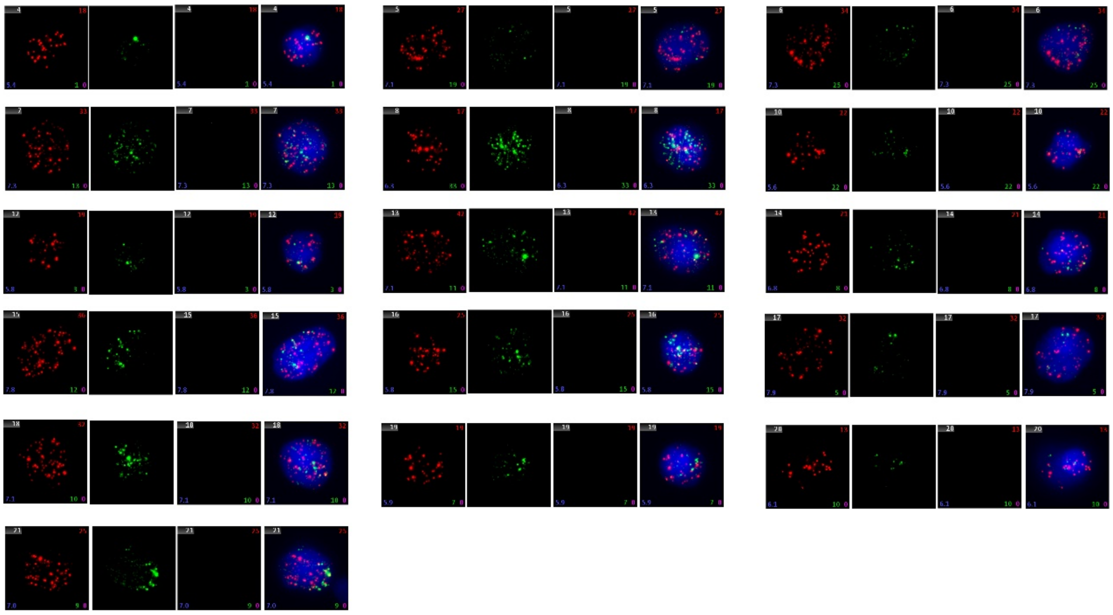

P8

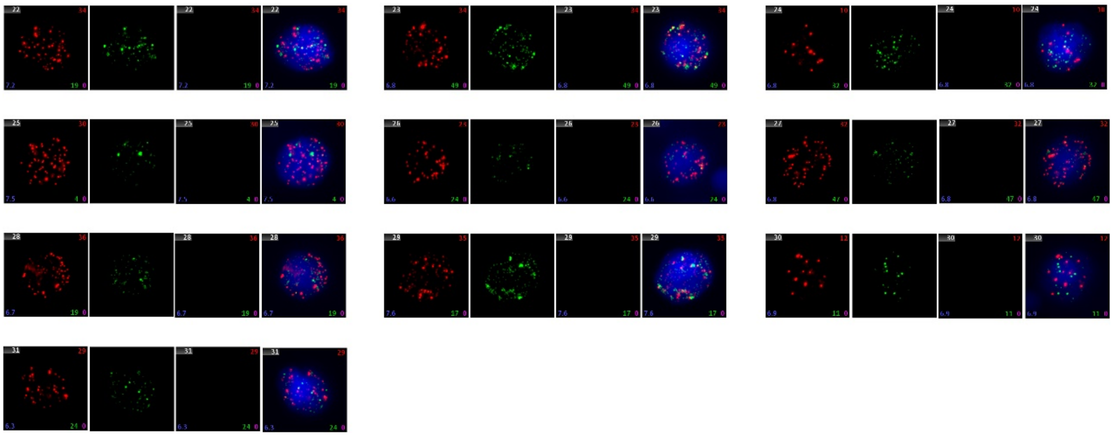

**Supplementary Figure 15**. Fluorescent pictures of the clinical CTCs captured by bio-orthogonal films and identified by the nucleus (blue), CD45 (white), EpCAM (red), and vimentin (green) staining (scale bar = 5 μm).

**P9**

*Bio-orthogonal film*

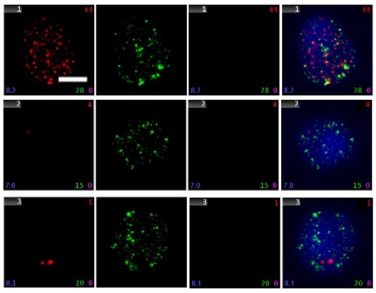

中科纳泰 nanopep | 肿瘤捕手 TUMORFISHER   *TumorFisher®*

外周血循环肿瘤细胞（CTC）检测结果报告

样本编号：SIAT202301　　送检日期：2023年8月16日

送检单位：暨南大学第二临床医学院

CTC检测结果

血液样本中富集和检测到符合 CTC 特征的细胞数：0 个。

① 荧光显微图像

（1）CTC：

| 合成图 | 明　场 | DAPI | CD45 | CK | CD45+DAPI | CK+DAPI |
|---|---|---|---|---|---|---|

CTC图片从检测指标中获取，最多展示2组照片。

（2）WBC：

| 合成图 | 明　场 | DAPI | CD45 | CK | CD45+DAPI | CK+DAPI |
|---|---|---|---|---|---|---|

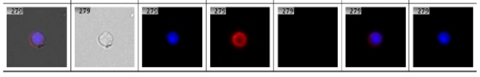

检测时间：2023年8月16日

**P10**

*Bio-orthogonal film*

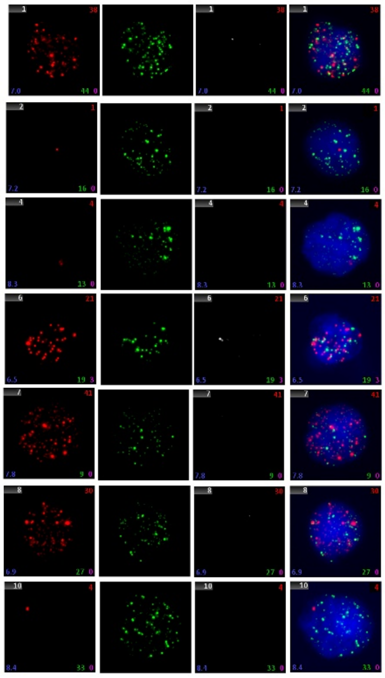

中科纳泰 nanopep | 肿瘤捕手 TUMORFISHER   *TumorFisher®*

外周血循环肿瘤细胞（CTC）检测结果报告

样本编号：SIAT202302　　送检日期：2023年8月24日

送检单位：暨南大学第二临床医学院

CTC检测结果

血液样本中富集和检测到符合 CTC 特征的细胞数：2 个。

① 荧光显微图像

（1）CTC：

| 合成图 | 明　场 | DAPI | CD45 | CK | CD45+DAPI | CK+DAPI |
|---|---|---|---|---|---|---|

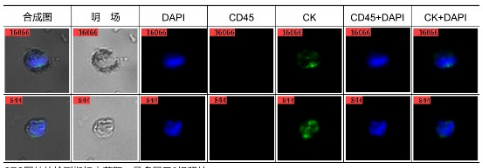

CTC图片从检测指标中获取，最多展示2组照片。

（2）WBC：

| 合成图 | 明　场 | DAPI | CD45 | CK | CD45+DAPI | CK+DAPI |
|---|---|---|---|---|---|---|

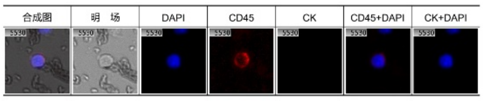

检测时间：2023年8月24日

**P11**

*Bio-orthogonal film*

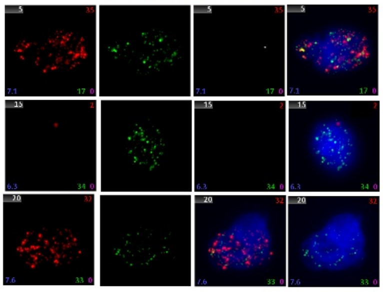

*TumorFisher®*

#### 外周血循环肿瘤细胞(CTC)检测结果报告

样本编号：SIAT202303　　　　送检日期：2023年9月4日

送检单位：暨南大学第二临床医学院

**CTC检测结果**

血液样本中富集和检测到符合 CTC 特征的细胞数： 1 个。

**1 荧光显微图像**

（1）CTC：

| 合成图 | 明　场 | DAPI | CD45 | CK | CD45+DAPI | CK+DAPI |
|---|---|---|---|---|---|---|

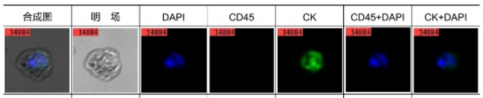

CTC图片从检测指标中获取，最多展示2组照片。

（2）WBC：

| 合成图 | 明　场 | DAPI | CD45 | CK | CD45+DAPI | CK+DAPI |
|---|---|---|---|---|---|---|

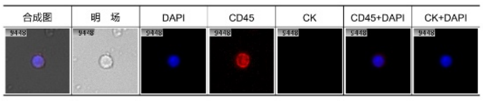

检测时间：2023年9月4日

**P12**

*Bio-orthogonal film*

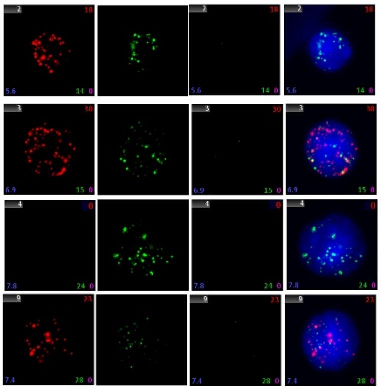

*TumorFisher®*

#### 外周血循环肿瘤细胞(CTC)检测结果报告

样本编号：SIAT202304　　　　送检日期：2023年9月13日

送检单位：暨南大学第二临床医学院

**CTC检测结果**

血液样本中富集和检测到符合 CTC 特征的细胞数： 3 个。

**1 荧光显微图像**

（1）CTC：

| 合成图 | 明　场 | DAPI | CD45 | CK | CD45+DAPI | CK+DAPI |
|---|---|---|---|---|---|---|

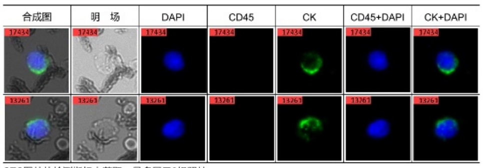

CTC图片从检测指标中获取，最多展示2组照片。

（2）WBC：

| 合成图 | 明　场 | DAPI | CD45 | CK | CD45+DAPI | CK+DAPI |
|---|---|---|---|---|---|---|

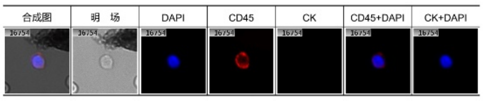

检测时间：2023年9月13日

**P13**

***Bio-orthogonal film***

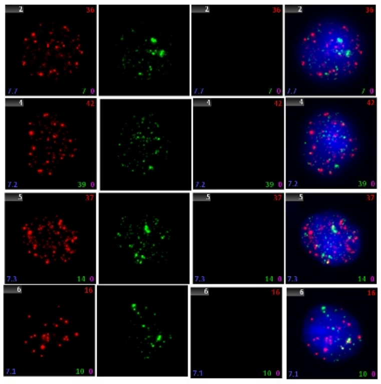

TumorFisher®

#### 外周血循环肿瘤细胞(CTC)检测结果报告

样本编号：SIAT202305　　送检日期：2023年9月23日

送检单位：暨南大学第二临床医学院

##### CTC检测结果

血液样本中富集和检测到符合 CTC 特征的细胞数： 2 个。

##### 1 荧光显微图像

（1）CTC：

| 合成图 | 明 场 | DAPI | CD45 | CK | CD45+DAPI | CK+DAPI |
|---|---|---|---|---|---|---|

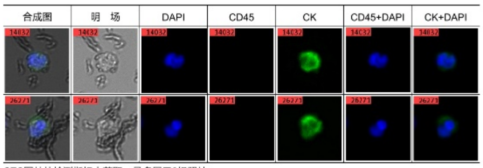

CTC图片从检测指标中获取，最多展示2组照片。

（2）WBC：

| 合成图 | 明 场 | DAPI | CD45 | CK | CD45+DAPI | CK+DAPI |
|---|---|---|---|---|---|---|

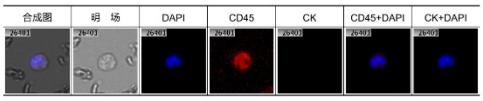

检测时间：2023年9月23日

**P14**

***Bio-orthogonal film***

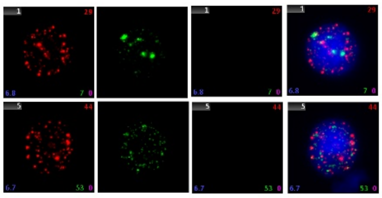

TumorFisher®

#### 外周血循环肿瘤细胞(CTC)检测结果报告

样本编号：SIAT202306　　送检日期：2023年9月23日

送检单位：暨南大学第二临床医学院

##### CTC检测结果

血液样本中富集和检测到符合 CTC 特征的细胞数： 2 个。

##### 1 荧光显微图像

（1）CTC：

| 合成图 | 明 场 | DAPI | CD45 | CK | CD45+DAPI | CK+DAPI |
|---|---|---|---|---|---|---|

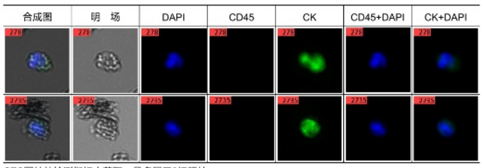

CTC图片从检测指标中获取，最多展示2组照片。

（2）WBC：

| 合成图 | 明 场 | DAPI | CD45 | CK | CD45+DAPI | CK+DAPI |
|---|---|---|---|---|---|---|

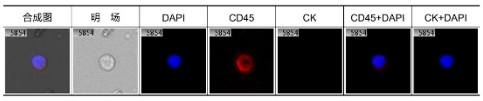

检测时间：2023年9月23日

**P15**

**Bio-orthogonal film**

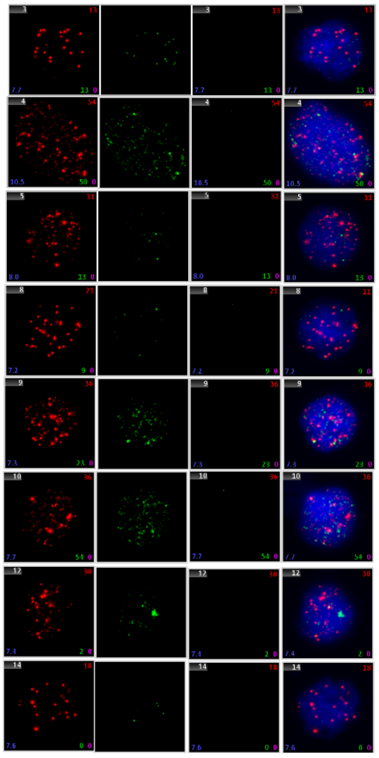

**TumorFisher®**

#### 外周血循环肿瘤细胞（CTC）检测结果报告

样本编号：SIAT202308　　送检日期：2023年9月23日

送检单位：暨南大学第二临床医学院

##### CTC检测结果

2mL 血液样本中富集和检测到符合 CTC 特征的细胞数：7 个。

###### 1 荧光显微图像

（1）CTC：

| 合成图 | 明 场 | DAPI | CD45 | CK | CD45+DAPI | CK+DAPI |
|---|---|---|---|---|---|---|

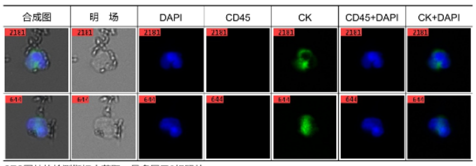

CTC图片从检测指标中获取，最多展示2组照片。

（2）WBC：

| 合成图 | 明 场 | DAPI | CD45 | CK | CD45+DAPI | CK+DAPI |
|---|---|---|---|---|---|---|

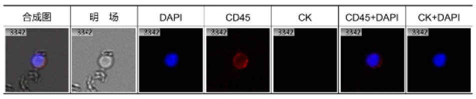

检测时间：2023年9月23日　　检测员：薛建

**P16**

**Bio-orthogonal film**

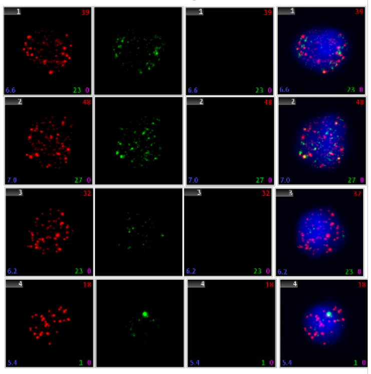

**TumorFisher®**

#### 外周血循环肿瘤细胞（CTC）检测结果报告

样本编号：SIAT202309　　送检日期：2023年9月27日

送检单位：暨南大学第二临床医学院

##### CTC检测结果

2mL 血液样本中富集和检测到符合 CTC 特征的细胞数：1 个。

###### 1 荧光显微图像

（1）CTC：

| 合成图 | 明 场 | DAPI | CD45 | CK | CD45+DAPI | CK+DAPI |
|---|---|---|---|---|---|---|

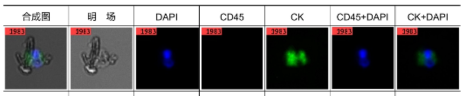

CTC图片从检测指标中获取，最多展示2组照片。

（2）WBC：

| 合成图 | 明 场 | DAPI | CD45 | CK | CD45+DAPI | CK+DAPI |
|---|---|---|---|---|---|---|

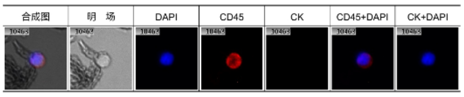

检测时间：2023年9月27日　　检测员：薛建

## P17

**Bio-orthogonal film**

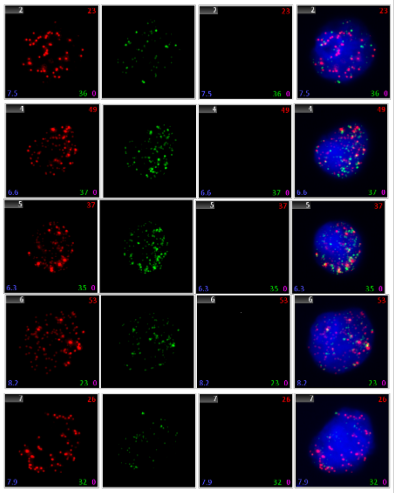

TumorFisher®

##### 外周血循环肿瘤细胞(CTC)检测结果报告

样本编号：SIAT202310　　　　送检日期：2023年9月27日

送检单位：暨南大学第二临床医学院

**CTC检测结果**

2mL 血液样本中富集和检测到符合 CTC 特征的细胞数：　1　个。

**1 荧光显微图像**

（1）CTC：

| 合成图 | 明 场 | DAPI | CD45 | CK | CD45+DAPI | CK+DAPI |
|---|---|---|---|---|---|---|

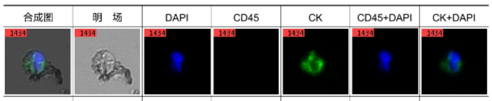

CTC图片从检测指标中获取，最多展示2组照片。

（2）WBC：

| 合成图 | 明 场 | DAPI | CD45 | CK | CD45+DAPI | CK+DAPI |
|---|---|---|---|---|---|---|

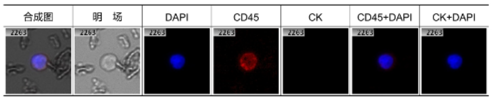

检测时间：2023年9月27日　　　　检测员：薛建

## P18

**Bio-orthogonal film**

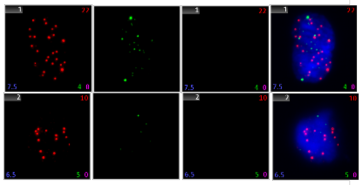

TumorFisher®

##### 外周血循环肿瘤细胞(CTC)检测结果报告

样本编号：SIAT202311　　　　送检日期：2023年9月27日

送检单位：暨南大学第二临床医学院

**CTC检测结果**

2mL 血液样本中富集和检测到符合 CTC 特征的细胞数：　2　个。

**1 荧光显微图像**

（1）CTC：

| 合成图 | 明 场 | DAPI | CD45 | CK | CD45+DAPI | CK+DAPI |
|---|---|---|---|---|---|---|

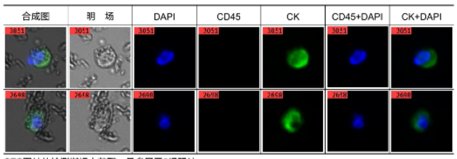

CTC图片从检测指标中获取，最多展示2组照片。

（2）WBC：

| 合成图 | 明 场 | DAPI | CD45 | CK | CD45+DAPI | CK+DAPI |
|---|---|---|---|---|---|---|

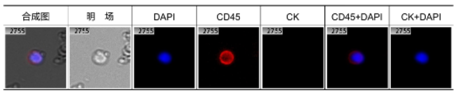

检测时间：2023年9月27日　　　　检测员：薛建

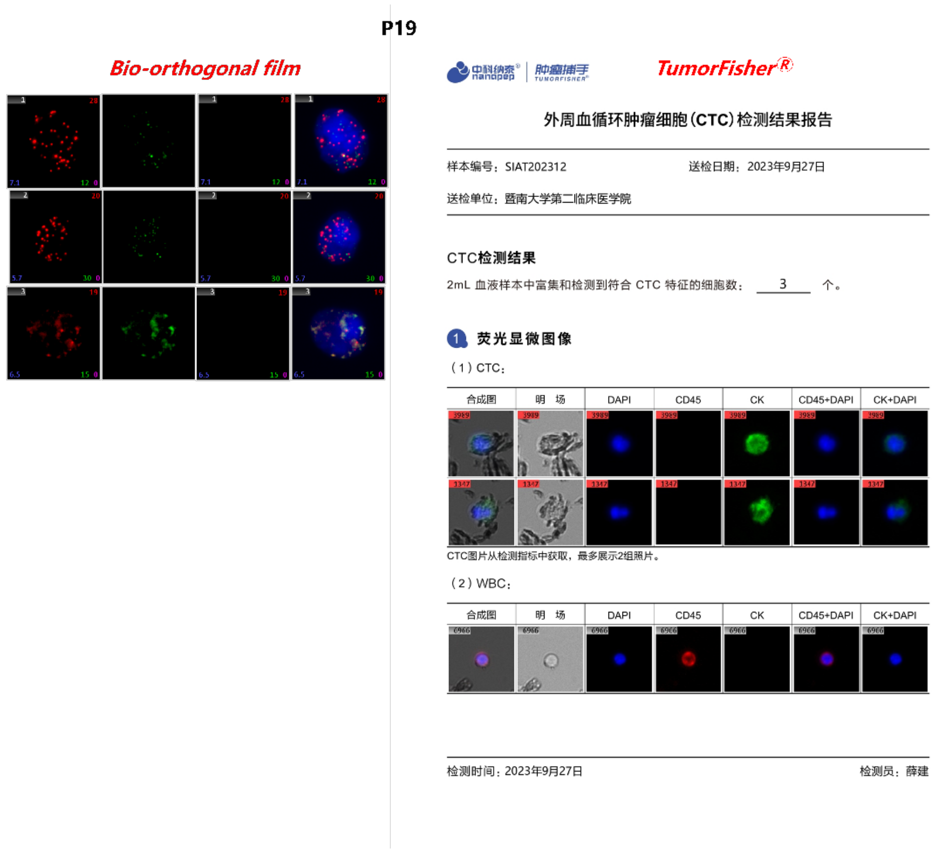

**Supplementary Figure 16**. Direct comparison between the bio-orthogonal films and TumorFisher® system about clinical CTCs detection. The CTCs captured by bio-orthogonal films are identified by the nucleus (blue), CD45 (white), EpCAM (red), and vimentin (green) staining, while the CTCs captured by TumorFisher® are identified by the nucleus (blue), CD45 (red) and CK (green) staining.

## P20

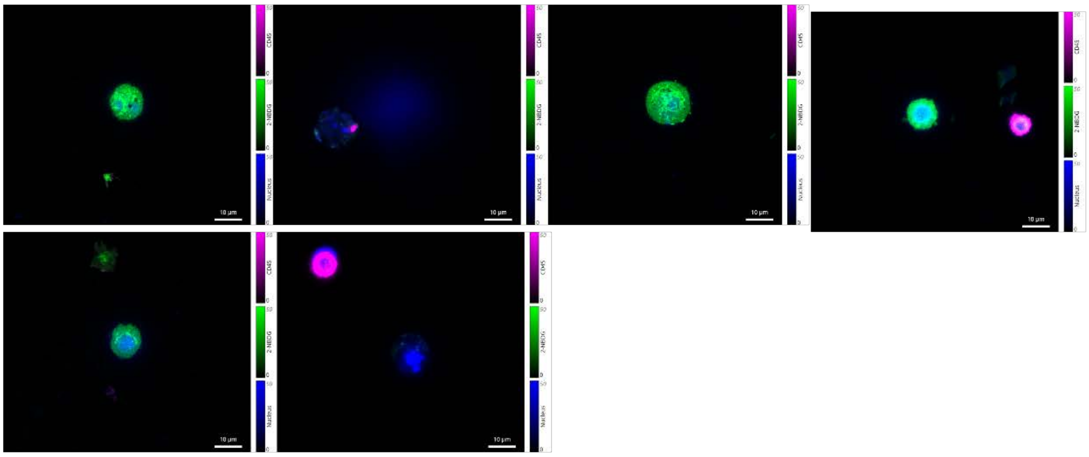

## P21

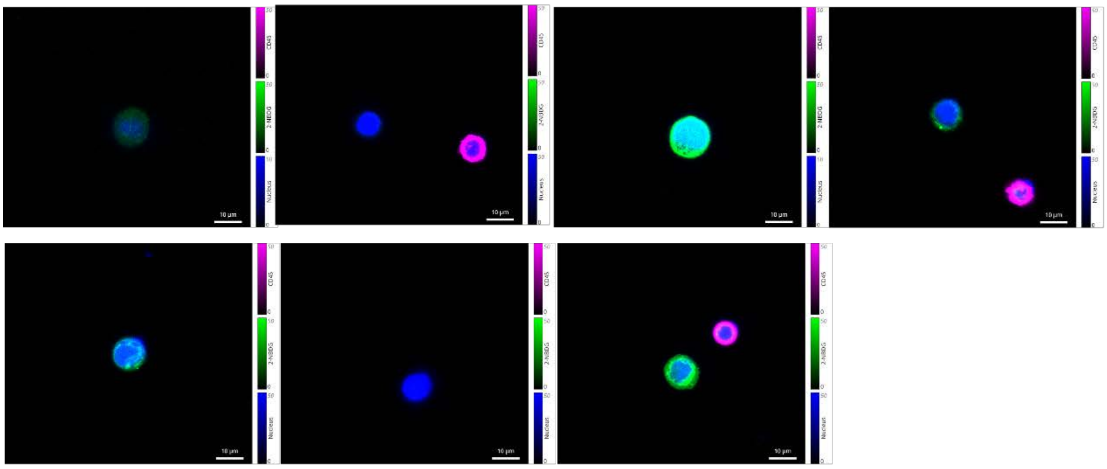

## P22

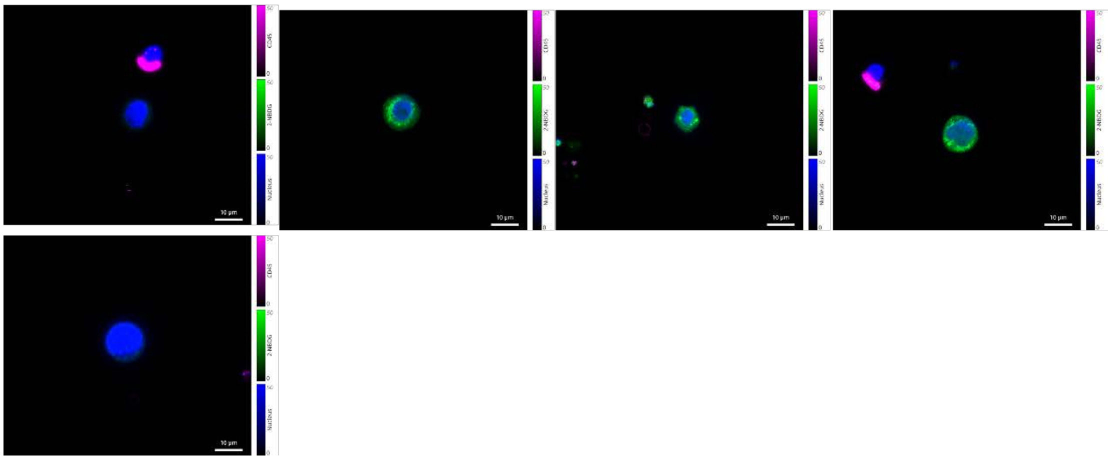

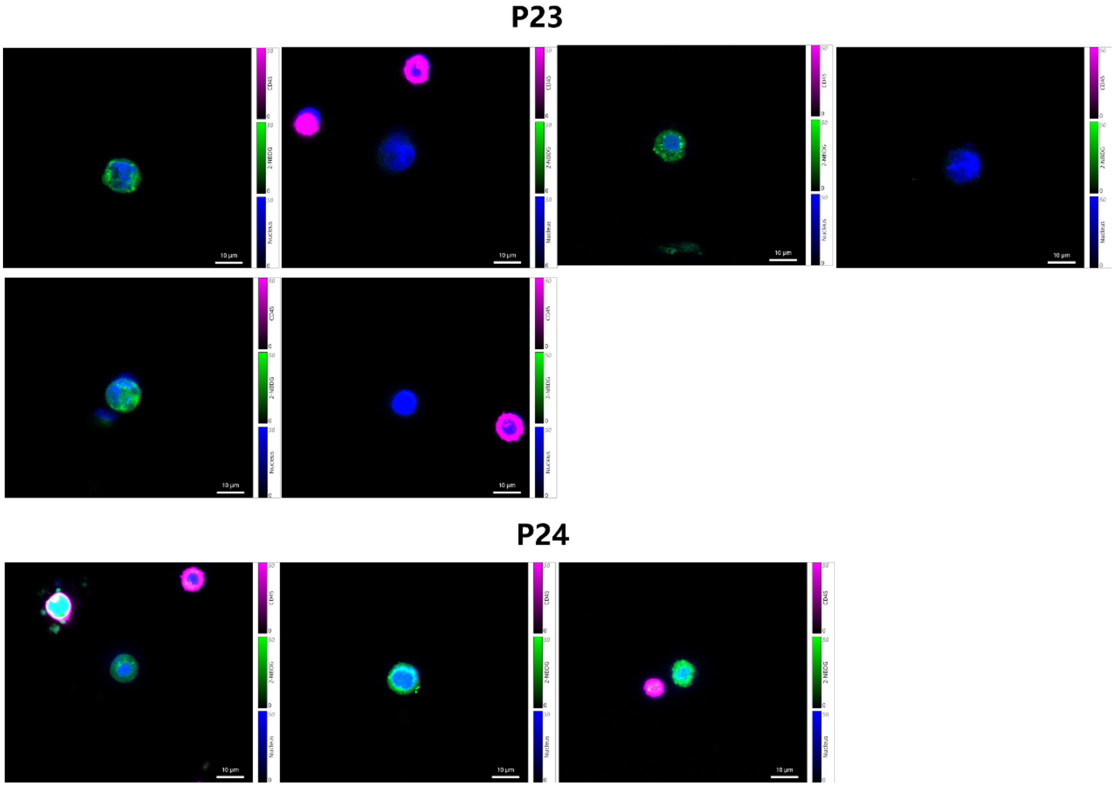

**Supplementary Figure 17**. Fluorescent pictures of the clinical CTCs captured by the bio-orthogonal films and released after the DTT treatment. The released CTCs are identified by the nucleus (blue) and CD45 (pink) staining, and the glycolytic activity of CTCs is examined by 2-NBDG (green) staining.

### Blank

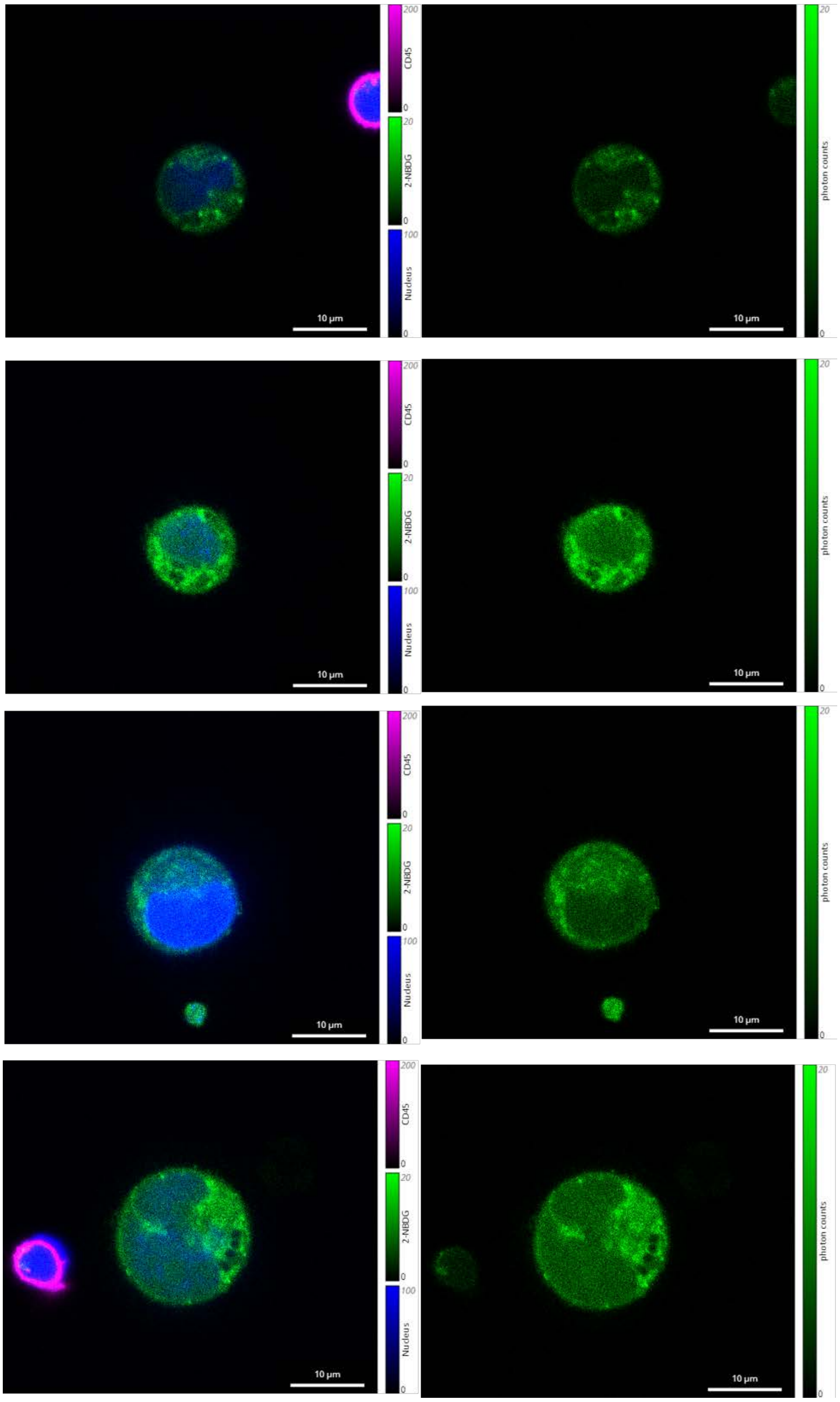

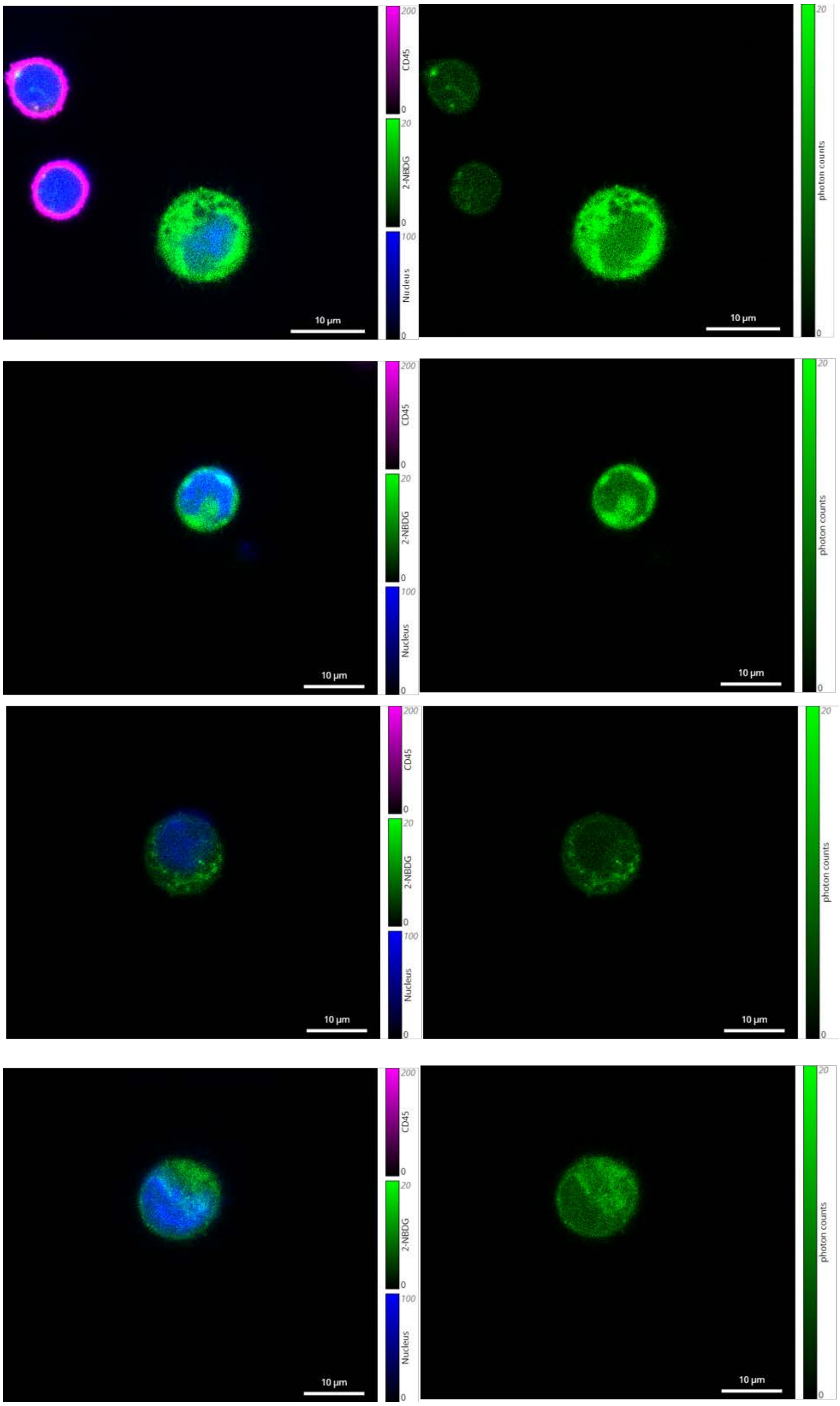
200
CD45
0
20
2-NBDG
0
100
Nucleus
0
10 µm
20
photon counts
0

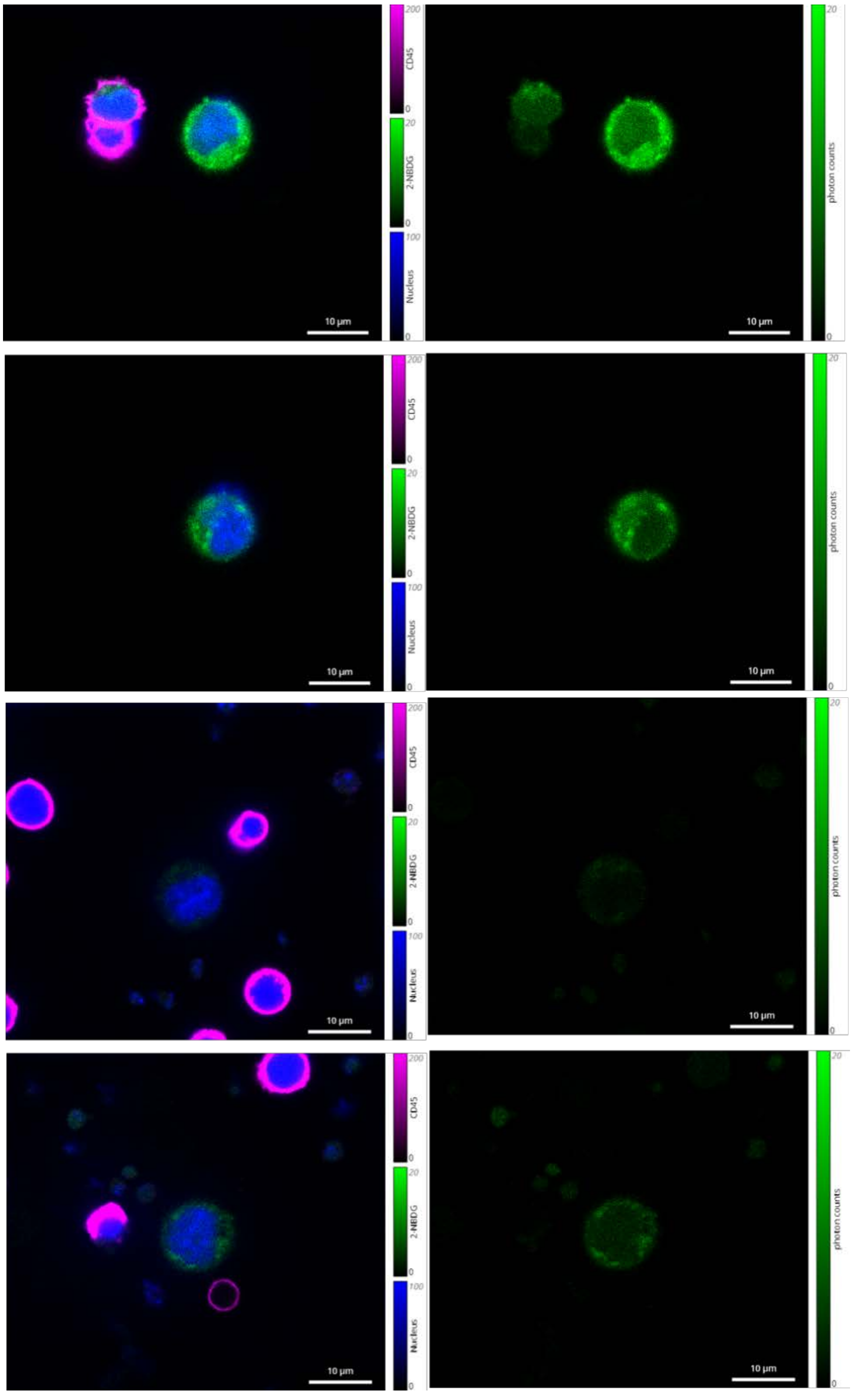
CD45
2-NBDG
Nucleus
photon counts
10 µm

**Fluorouracil**

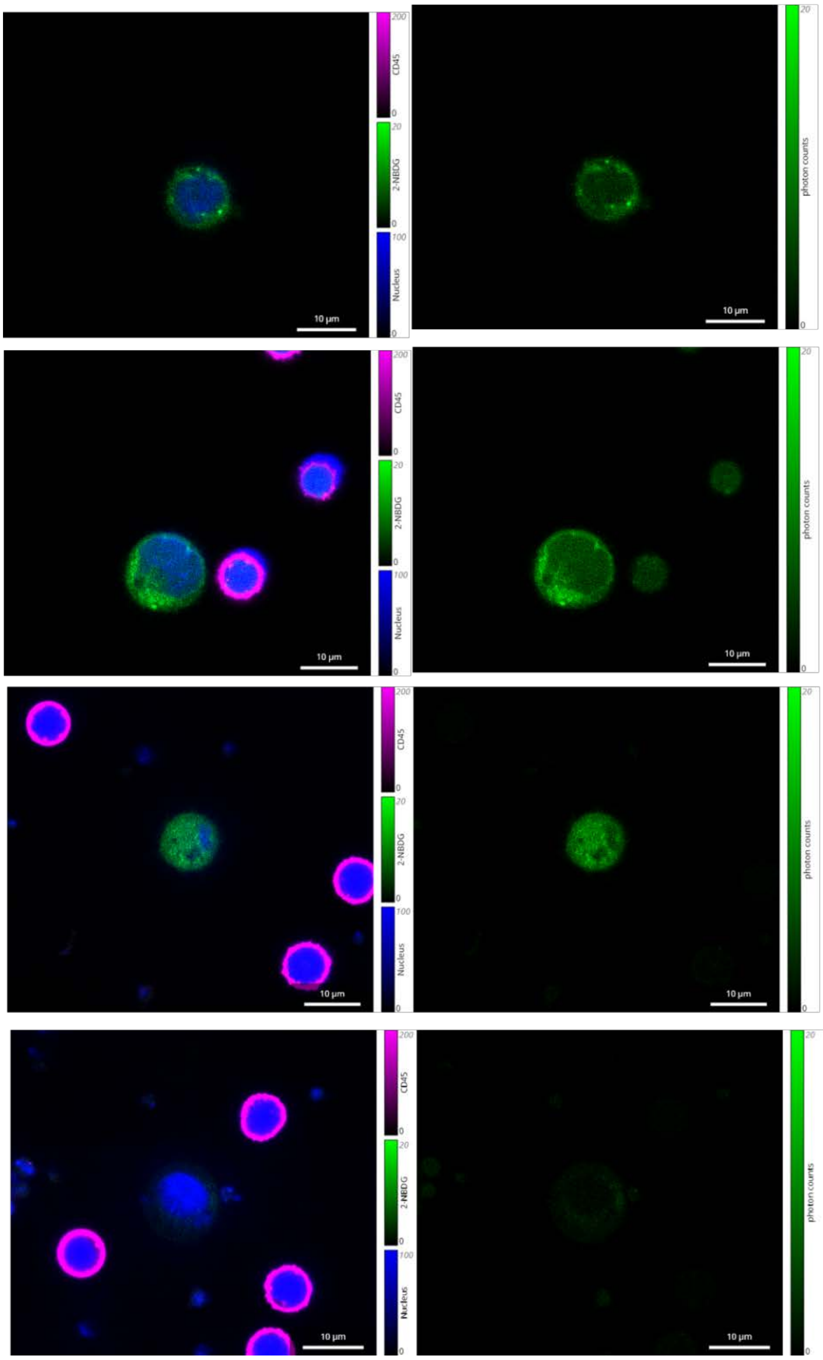

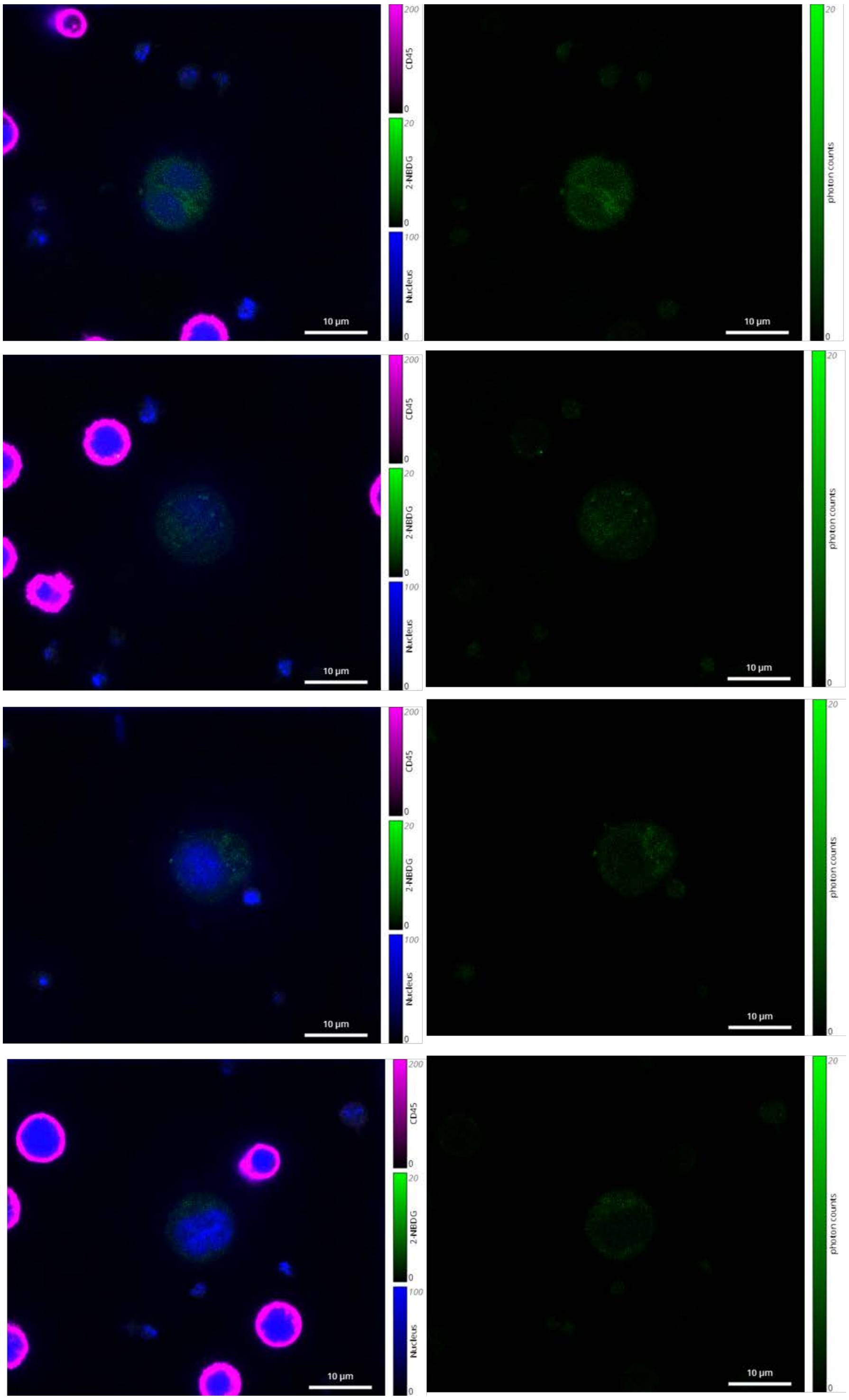
CD45
2-NBDG
Nucleus
photon counts
10 μm

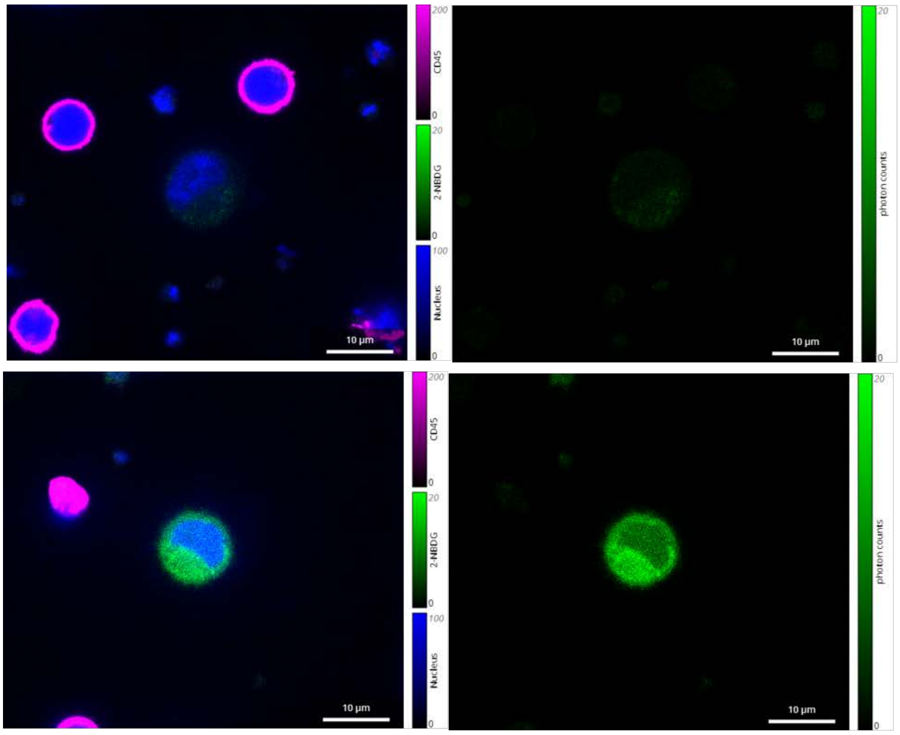

**Oxaliplatin**

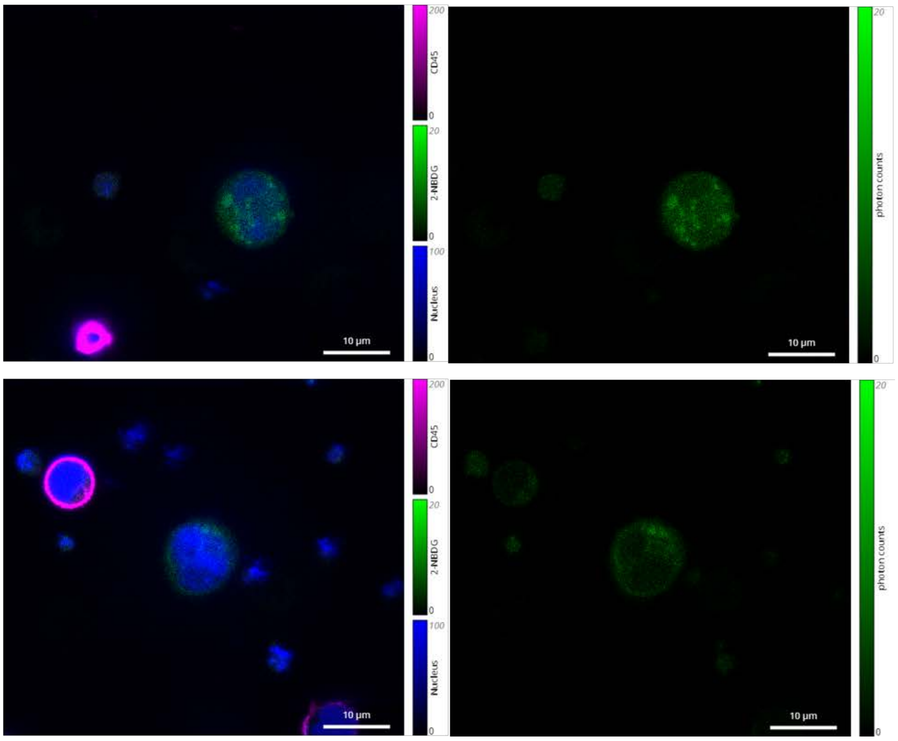

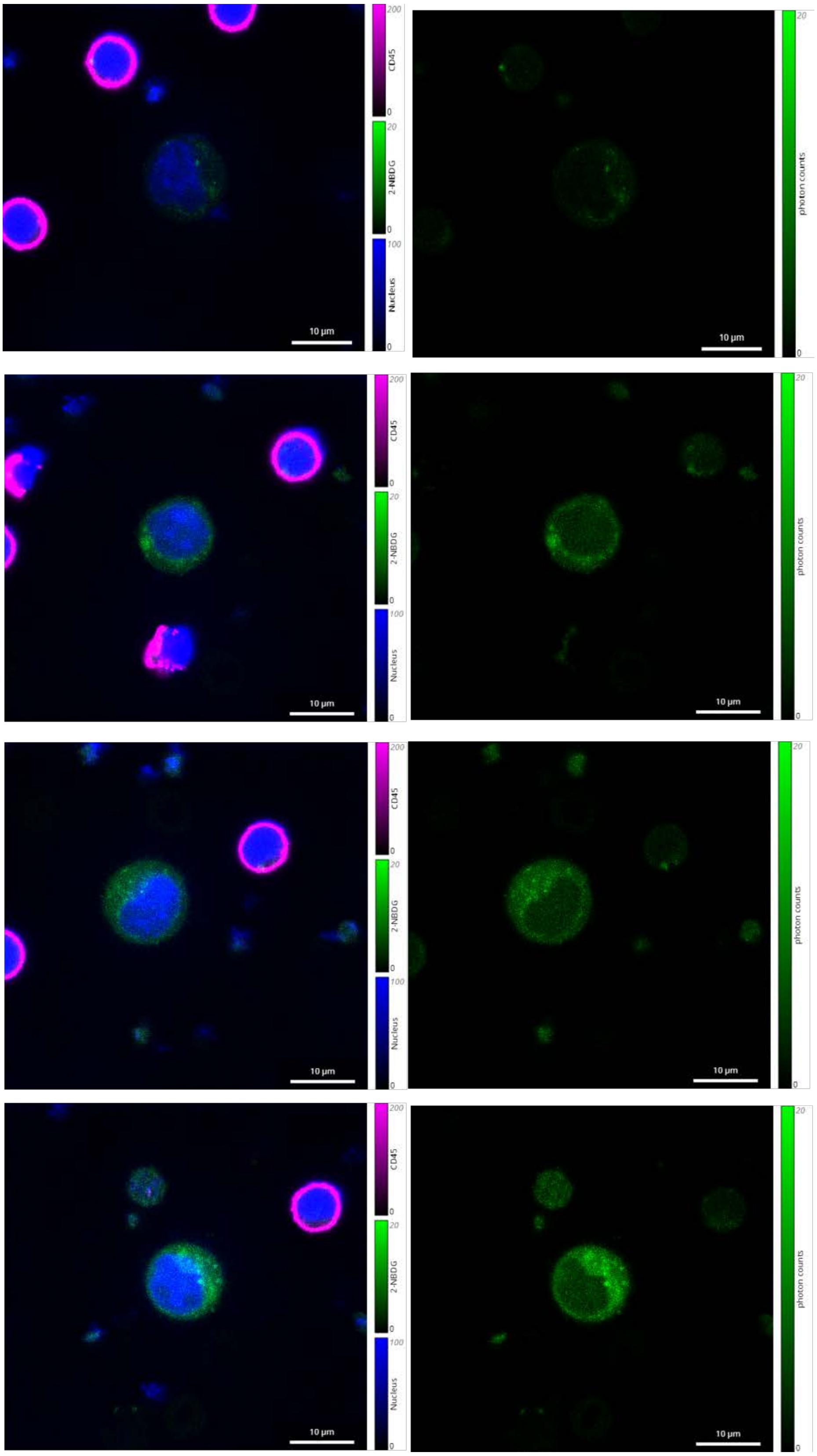

200
CD45
0
20
2-NBDG
0
100
Nucleus
0
10 µm
20
photon counts
0
10 µm
200
CD45
0
20
2-NBDG
0
100
Nucleus
0
10 µm
20
photon counts
0
10 µm
200
CD45
0
20
2-NBDG
0
100
Nucleus
0
10 µm
20
photon counts
0
10 µm
200
CD45
0
20
2-NBDG
0
100
Nucleus
0
10 µm
20
photon counts
0
10 µm

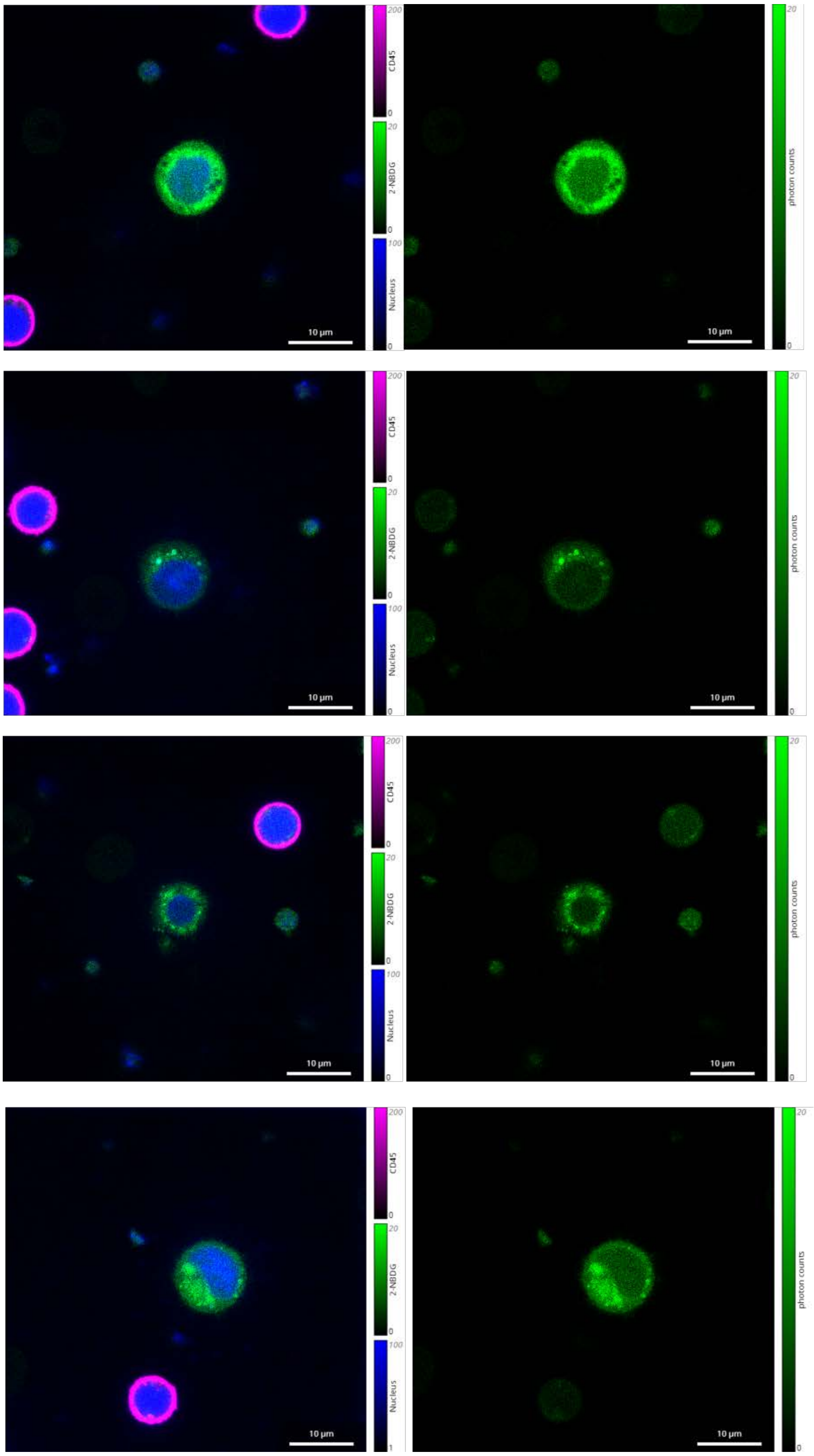

CD45
2-NBDG
Nucleus
photon counts
10 μm

**Supplementary Figure 18.** Fluorescent pictures of the released CTCs in the drug susceptibility test (Fluorouracil and Oxaliplatin). The released CTCs are identified by the nucleus (blue) and CD45 (pink) staining, and the glycolytic activity of CTCs in different groups is examined by 2-NBDG (green) staining.

**Supplementary Table 1.** Information of donors.

| No. | Type | Gender | Age |
|---|---|---|---|
| **H1** | healthy | M | 36 |
| **H2** | healthy | F | 67 |
| **H3** | healthy | M | 34 |
| **H4** | healthy | F | 32 |
| **P1** | breast cancer | F | 67 |
| **P2** | liver cancer | M | 62 |
| **P3** | liver cancer | M | 57 |
| **P4** | breast cancer | F | 59 |
| **P5** | gastric cancer | F | 65 |
| **P6** | nasopharyngeal carcinoma | M | 57 |
| **P7** | lung cancer | M | 54 |
| **P8** | liver cancer | F | 49 |
| **P9** | colon cancer (partly mesenchymal tissue tumor) | M | 28 |
| **P10** | rectal cancer (mesenchymal tissue tumor) | M | 59 |
| **P11** | rectal cancer (mesenchymal tissue tumor) | M | 70 |
| **P12** | osteosarcoma | M | 14 |
| **P13** | cervical cancer | F | 37 |
| **P14** | cervical cancer | F | 64 |
| **P15** | rectal cancer | M | 68 |

| | | | |
|---|---|---|---|
| **P16** | nasopharyngeal carcinoma | M | 22 |
| **P17** | nasopharyngeal carcinoma | M | 56 |
| **P18** | cervical cancer | F | 58 |
| **P19** | colon cancer (Stage III) | M | 51 |
| **P20** | colon cancer (Stage III) | M | 55 |
| **P21** | colon cancer (Stage III) | M | 62 |
| **P22** | colon cancer (Stage III) | F | 45 |
| **P23** | colon cancer (Stage III) | M | 61 |
| **P24** | colon cancer (Stage III) | F | 52 |
| **P25** | colon cancer (Stage III) | M | 58 |

**Supplementary Table 2.** Comparison of WBCs contamination between our technique and other reported clinical CTCs detection methods.

| | Cancer Type | Methods | WBCs |
|---|---|---|---|
| **Ref. 1** | Prostate cancer | EpCAM targeting | 58-9249 per mL |
| **Ref. 2** | Non-small cell lung cancer | EpCAM targeting | 326 ± 165 per mL |
| **Ref. 3** | Breast cancer | EpCAM targeting | 350.7 ± 122.8 per 5 mL |
| **Ref. 4** | Breast cancer | Physical filtration | 663 ± 647 per mL |
| **Our study** | Covering 10 cancer types | MGE | 341-2750 per mL |
